# Integrative proteomic analysis and molecular dynamics simulations of ANKLE2 reveal mechanisms of microcephaly

**DOI:** 10.64898/2026.09.28.755110

**Authors:** Adam T. Fishburn, Vardhan Peddamallu, Nicholas J. Lopez, Ines M. Bilkic, Isabella J. Hixson, Edwina Mambou, Heng Pan, Boris Bonaventure, Cole J. Florio, Chase L. S. Skawinski, Michelle Lopez Ramos, Parth R. Bandivadekar, Sydney S. Becker, Bhargavi L. Gidugu, Judith L. A. Fishburn, Alana P. Gudinas, Jacqueline H. Barlow, Danielle J. Mai, Surl-Hee Ahn, Jeffrey R. Johnson, Nichole Link, Priya S. Shah

## Abstract

ANKLE2 is a scaffolding protein with crucial roles in neuroprogenitor cell division and embryonic brain development. Pathogenic variants in *ANKLE2* cause primary microcephaly, a congenital disorder characterized by impaired neurodevelopment. Nonetheless, the underlying dysfunction of ANKLE2 is not understood. Here, we define the ANKLE2 protein interaction landscape, which includes cell division proteins and novel microcephaly candidates in the APC (anaphase-promoting complex). Analysis of six pathogenic variants reveals changes in this interactome that likely result in microcephaly. Using molecular simulations, we identify the structural consequences of five pathogenic substitutions, including disruption of important helical structures. Combined, these techniques suggest loss of interaction with the PP2A (protein phosphatase 2A) complex is a common mechanism for ANKLE2 pathogenesis. Phosphoproteomics reveals widespread alterations in ANKLE2- and PP2A-dependent phosphorylation of cell division proteins. Together, our integrative proteomics and simulation approach improves the molecular understanding of ANKLE2 function and its pathogenic changes in primary microcephaly.

**Highlights:**

- ANKLE2 interacts with the APC (anaphase-promoting complex), whose loss results in microcephaly *in vivo*.
- Pathogenic ANKLE2 variants have distinct interaction networks and structural dynamics.
- Mutations in Caulimovirus Domain disrupt domain structure and interaction with PP2A (protein phosphatase 2A) complex.
- ANKLE2 regulates phosphorylation of many proteins through PP2A, including APC.

## Introduction

The scaffolding protein ANKLE2 (ankyrin repeat LEM domain-containing 2) has emerged as a key regulator of cell division and brain development ^1^. ANKLE2 regulates nuclear envelope disassembly and reassembly dynamics by coordinating the phosphorylation state of BAF (barrier to autointegration factor) through interactions with the kinase VRK1 (vaccinia related kinase 1) and the PP2A (protein phosphatase 2A) complex ^2^. Depletion of the *ANKLE2* homolog in *C. elegans* (*lem-4L*) results in abnormal nuclear morphology driven by increases in BAF phosphorylation, leading to embryonic lethality ^3^. A genetic screen in *Drosophila* first identified that mutation of the *ANKLE2* homolog (*Ankle2* in flies) results in abnormal neurodevelopment and reduced larval brain volume ^4^. Underlying this phenotype is a disruption of spindle pole alignment and cell division in neuroprogenitor cells ^5^. Importantly, introduction of human ANKLE2 can rescue these phenotypes in flies, supporting that ANKLE2’s function in brain development is conserved across these models ^5–7^.

Defects in neuroprogenitor cell division often result in primary microcephaly, a condition involving a significant reduction in brain and head size at birth (2-3 standard deviations below mean circumference for sex/age) ^8,9^. Microcephaly is commonly accompanied by a variety of clinical symptoms, including significant cognitive and physical developmental delays, sensory deficits, and seizures ^10–12^. Rare pathogenic variants in at least 31 genes, including those regulating spindle assembly, cause primary microcephaly (MCPH) ^8^. *ANKLE2* is one such MCPH gene. Study of unattributed human Mendelian diseases revealed mutations in *ANKLE2* (p.L573V/p.Q782*) were responsible for MCPH in two individuals ^4^. To date, 17 pathogenic variants of *ANKLE2* have been identified and cause ANKLE2-related MCPH, otherwise known as MCPH16 ^4–6,13,14^. Nearly all cases of MCPH16 are identified with congenital microcephaly apparent at birth or *in utero*, although in several cases microcephaly developed postnatally ^6,14^. Brain MRI of MCPH16 cases consistently reveal cortical dysplasia, extra axial spaces, and thinning or partial agenesis of the corpus callosum. Behavioral impacts include speech and language developmental delays and impacts to gross and fine motor function ^6^. Given the range of pathogenic variants and the fact that most MCPH16 cases are biallelic, it is challenging to relate specific mutations to pathogenesis.

The importance of ANKLE2 in human brain development is clear, but less is known about its molecular functions. One limitation is that the structure of ANKLE2 has not yet been resolved and is complicated by its many unstructured loops. Our previous work used predictive structural analysis to define six distinct structural elements within ANKLE2 ^15^. A N-terminal transmembrane (TM) domain anchors ANKLE2 to the ER membrane ^16^. Following the TM domain is the LEM domain (named after <u>L</u>AP2, <u>E</u>merin, and <u>M</u>AN1), which mediates binding to BAF at the nuclear lamina ^17^. The LEM domain is a ∼50 amino acid structure consisting of three α-helices in a conserved orientation ^15^. Next is the caulimovirus domain (CD), a unique ∼55 amino acid structure comprising a small β-sheet sandwiched by two α-helices. CD supports binding with the PP2A complex composed of PPP2R1A (scaffold), PPP2R2D (regulatory), and PPP2CB (catalytic) subunits, which together regulate dephosphorylation of BAF during nuclear envelope reassembly ^2,18^. Following the CD is the ankyrin repeat domain (from here on referred to as ANK). This domain is highly conserved among ANKLE2 orthologs and is thought to serve in ANKLE2’s scaffolding capacity. The C-terminal half of the protein contains two structured regions with high predicted structural conservation but poorly defined function ^15^. We previously referred to these as uncharacterized domains 1 and 2 (Unc1 and Unc2). Unc1 (amino acids 474-604) resembles GIY-YIG nuclease domains, similar to ANKLE1, but lacks catalytic endonuclease activity ^15^. Despite the elusive role of Unc1 and Unc2, we speculate they serve critical functions since they exist in nearly all orthologs ^15^. Consistent with this, nonsense mutations that ablate these structures are pathogenic in humans ^4,5^ and do not rescue brain development in mutant flies ^5,7^.

All 17 known pathogenic variants of *ANKLE2* either introduce missense mutations within structured domains or splicing/nonsense mutations that abolish them, underscoring the functional importance of these domains (Figure 1A). Intrinsically disordered regions (IDRs) can mediate protein interactions ^19–21^, yet benign variants occur primarily in IDRs (gnomAD) ^22^. Taken together, we hypothesized that pathogenic mutations disrupt structured domains to perturb ANKLE2’s scaffolding functions, ultimately cascading to microcephaly. To test this hypothesis, we used an integrative proteomic and molecular dynamics (MD) simulation approach (Figure 1B). We defined the ANKLE2 protein interaction network and changes associated with six pathogenic variants using affinity purification and mass spectrometry (AP-MS). This comprehensive analysis revealed new links between ANKLE2, mitochondrial proteins, and the APC (anaphase-promoting complex). Loss of *ANKLE2* governs both mitochondrial protein abundance and activity. Knockdown of APC members in developing flies revealed significant reduction in brain volumes, suggesting APC members are candidate MCPH genes. Pathogenic variants lost key interactions with PP2A and APC. In parallel, we characterized the structural dynamics of ANKLE2 and associated changes for five out of six of these pathogenic variants using Gaussian accelerated molecular dynamics (GaMD) simulations. These simulations serve as a “computational microscope” with atomistic resolution. Our GaMD simulations revealed both dramatic and subtle changes in ANKLE2 secondary structure, which we speculate drive similarly dramatic and subtle changes in ANKLE2 protein interactions. Insights from proteomics and GaMD for variants which lost PP2A interactions led us to perform global phosphoproteomics, revealing how ANKLE2-dependent PP2A phosphatase activity regulates much more than BAF. Thus, by combining proteomics and GaMD, we move from systems to structure and back again, gaining molecular understanding of how pathogenic ANKLE2 variants cause disease.

**Figure 1:**
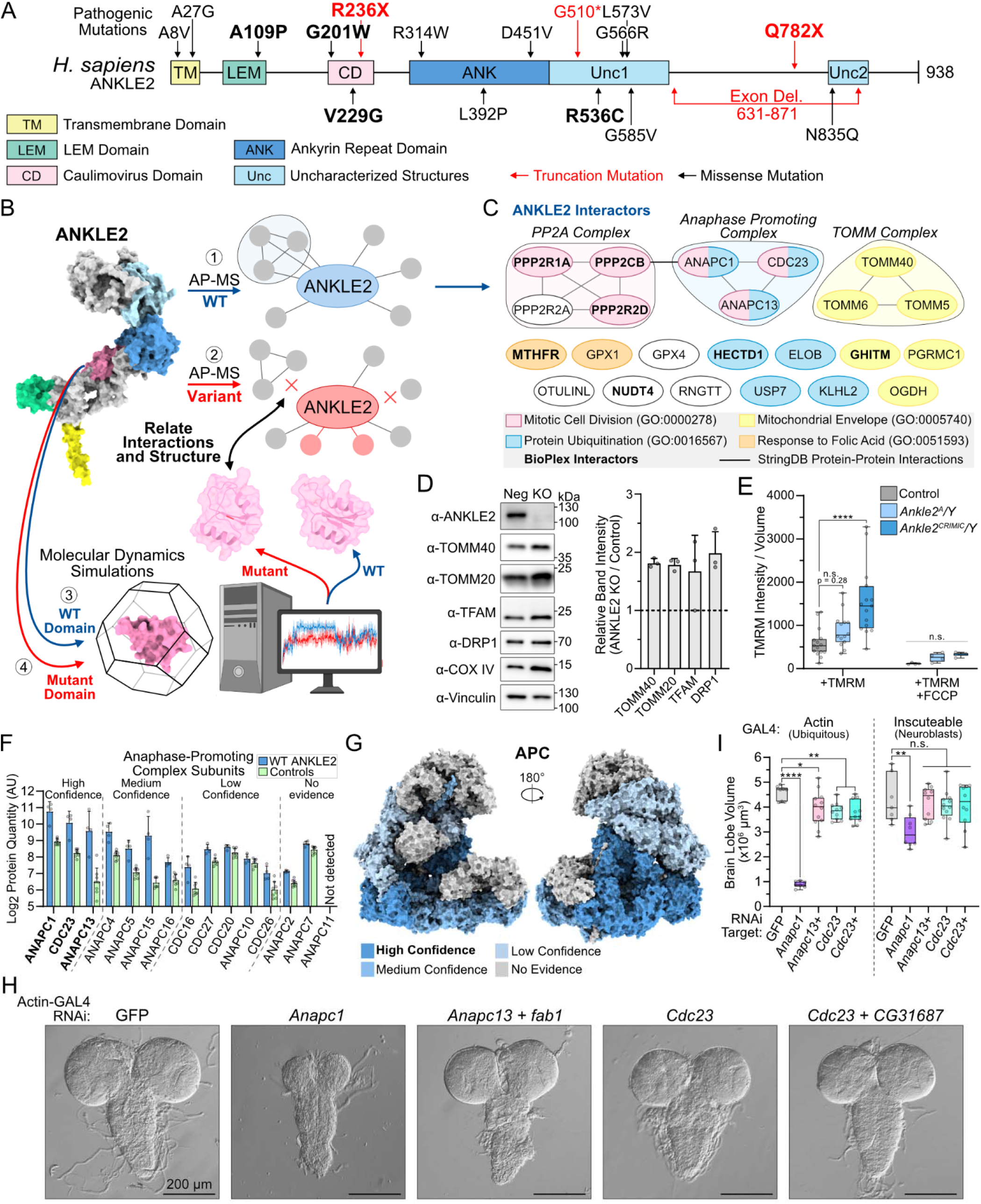
Mapping ANKLE2 protein interaction reveals interactions with mitochondrial proteins and APC. **A)** Diagram of ANKLE2 with structured domains annotated as previously described ^15^ with known pathogenic variants associated with congenital microcephaly. **B)** Schematic overview of integrative proteomics and molecular dynamics approach. **C)** Visualization of ANKLE2 protein interaction network. **D)** Western blot of Huh7 ANKLE2 KO cells to determine changes in mitochondrial protein abundance. TOMM complex function was assessed by attempting to visualize un-processed TFAM (expected band at ∼29 kDa). Quantification of western blot band intensities from three independent replicates. Signal was first normalized to vinculin loading control and then set relative to control cells. **E)** Box plot of mitochondrial membrane potential of third instar larval flies, measured by TMRM. Gray dots indicate individual animals, 5-6 animals were imaged per experiment, and three independent experiments were performed. One-way ANOVA with Dunnett’s multiple comparisons test, n.s. = p > 0.05, **** = p < 0.0001. **F)** Proteomics protein quantity measurements for ANKLE2 and all APC subunits. Confidence level indicative of number of criteria met (high = 4/4, medium = 3 or 2/4, low = 1/4, no evidence = 0/4). **G)** Confidence of ANKLE2 interaction overlayed onto the APC structure (pdb: 5A31) ^36^. **H)** Bright field images of third-instar larval brains from ubiquitous knockdown (*Act-GAL4*) using *in vivo* RNAi of the indicated APC subunits. **I)** Quantification of brain volume from a single brain lobe with ubiquitous knockdown (left, *Act-GAL4*) and neural stem cell-specific knockdown (right, *inscuteable-GAL4*). Gray dots indicate individual animals, n = 7-12. One-way ANOVA with Dunnett’s multiple comparisons test. * p < 0.05, ** p <0.01, **** p < 0.0001, n.s. not significant (p > 0.05).

## Results

### Defining the ANKLE2 protein interaction landscape

To better understand the molecular role of ANKLE2, we first sought to define the extent of its protein interaction landscape. FLAG-tagged ANKLE2 was expressed in HEK293T cells, which were used to maximize protein expression and capture a wide breadth of interacting proteins. These cells are commonly used for proteomic studies, including those focused on neurological diseases ^23^. To capture and identify interacting proteins, we used affinity purification and mass spectrometry (AP-MS) (Figure S1A-C). Since ANKLE2 is ER-localized, we employed both FLAG-tagged GFP and ER-localized GFP (ER-GFP) as controls to account for organelle-specific background. Protein and peptide MS data for ANKLE2 interactions were compared to controls and analyzed using a combination of statistical techniques (Figure S1D-F) ^24,25^. This extensive filtering of ANKLE2-protein interactions yielded 23 high-confidence protein-protein interactions (HC-PPIs) for ANKLE2 (Figure 1C, Supplementary Data 1). Our interactome was significantly enriched for known ANKLE2 interactors from BioPlex 3.0 ^26^ (Fisher’s exact, p = 6.69 x 10^-11^), suggesting our approach captured biologically relevant interactions.

To gain insight into the high-level processes involved with ANKLE2-interactors, we visualized HC-PPI interconnections using STRINGdb ^27^ and performed functional enrichment analysis using g:GOSt ^28^. We observed the expected enrichment for the PP2A complex and cell division biological processes (Figure S1G). We also identified enrichment for other processes, especially those involved in outer mitochondrial membrane transport and ubiquitination. Neither of these processes have been associated with ANKLE2 function previously, prompting us to functionally investigate these novel interactions.

### Loss of *Ankle2* alters mitochondrial function *in vivo*

The identification of three proteins from the translocase of the outer mitochondrial membrane (TOMM) complex in our ANKLE2 interactome was unexpected (Figure 1C). The TOMM complex is present in nearly all eukaryotes and serves as a key site of mitochondrial protein import into the intermembrane space ^29^. We identified three of the seven human TOMM complex subunits as HC-PPIs (TOMM40, TOMM5, and TOMM6). TOMM40, a highly conserved β-barrel, acts as the central component onto which other receptor and accessory subunits (including TOMM5 and TOMM6) attach ^30^.

We speculated that ANKLE2 may play a role in TOMM complex protein import function from the cytosol. To test this, we evaluated the processivity of the protein TFAM (transcription factor A, mitochondrial) in Huh7 ANKLE2 knockout (KO) cells we generated previously ^31^. TFAM is a nuclear-encoded mitochondrial DNA polymerase that is produced outside the mitochondria in a 29 kDa form, and imported by TOMM, where it is then proteolytically cleaved into an active 25 kDa form ^32,33^. We observed only processed TFAM in both control and ANKLE2 KO cells, suggesting that ANKLE2 does not play a role in TOMM complex importing functions (Figure 1D). However, we did observe an increase in total TFAM abundance in ANKLE2 KO cells when compared to control cells. In fact, the abundance of multiple mitochondrial proteins (TOMM40, TOMM20, TFAM, DRP1, COX IV) were moderately increased in ANKLE2 KO cells, suggesting some indirect role of ANKLE2 in mitochondrial function (Figure 1D).

We next tested if loss of *Ankle2* alters mitochondrial function using an *in vivo* fruit fly model ^4,5,7^. We used two *Ankle2* loss-of-function (LOF) mutants for these studies. *Ankle2^A^*mutants contain a L326H mutation in the ANK domain ^4^ and have moderate brain development phenotypes ^5^. In contrast, *Ankle2^CRIMIC^* flies, which are CRISPR mutagenized to produce an early truncation of *Ankle2*, have more severe brain development phenotypes ^5^. Mitochondrial membrane potential was measured in larval muscle tissue from wild type (WT) and mutant *Ankle2* third instar larval flies using tetramethylrhodamine methyl ester (TMRM). We observed modest increases in TMRM signal in *Ankle2^A^*flies relative to WT, and a substantial and statistically significant increase in *Ankle2^CRIMIC^* flies (Figure 1E). Notably, this increase in mitochondrial membrane potential following loss of *Ankle2* correlates with degree of loss and severity of other developmental phenotypes ^5^. Together, we identify a novel protein interaction between ANKLE2 and the TOMM complex. ANKLE2 regulates mitochondrial protein abundance and membrane potential, and whose dysfunction may be related to pathogenesis caused by loss of ANKLE2.

### The Anaphase-Promoting Complex is an ANKLE2 interactor and microcephaly candidate

One of the most interesting HC-PPIs we identified was APC. APC is a large, E3-ubiquitin ligase complex which acts as a key regulator of the cell cycle by degrading proteins to mediate chromosome separation and mitotic exit ^34,35^. Of this complex we identified ANAPC1 (anaphase-promoting complex, subunit 1), ANAPC13 (anaphase-promoting complex, subunit 13), and CDC23 (cell division cycle protein 23 homolog, also known as ANAPC8) as HC-PPIs (Figure 1C). We also detected nearly every other subunit of this complex in our proteomic data (all except ANAPC11, Figure 1F). However, many of these complex members narrowly failed our strict criteria. When overlaid on the APC structure ^36^, we observed that interaction confidence was correlated with neighboring scaffolding subunits (ANAPC1, 4, 5, CDC23, etc., Figure 1F-G, S1H). This suggests that ANKLE2 binds APC via the scaffolding domain and regulates interactions between APC and its substrates, rather than being a target of its ubiquitin ligase activity.

To test if loss of APC causes microcephaly, we utilized a larval fruit fly model of brain development. Ubiquitous and tissue-specific RNAi knockdown was performed on the fly orthologs of APC subunits (*Anapc1, Anapc13,* and *Cdc23*) using a GAL4-UAS system ^37^. Ubiquitous knockdown of these genes resulted in decreased larval brain volume, with the most severe phenotypes occurring with *Anapc1* (Figure 1H-I). This phenotype was also present to a lesser extent in neural stem cell-specific knockdown (Figure 1I). Taken together, we conclude that ANKLE2 interacts with APC, whose LOF causes microcephaly in flies.

### ANKLE2 pathogenic variants disrupt interactions with proteins critical for brain development

Given ANKLE2’s scaffolding functions, we hypothesized that these ANKLE2 interactions are perturbed in instances of congenital microcephaly. We therefore sought to profile the protein-protein interactions for various ANKLE2 pathogenic variants. Mutations throughout *ANKLE2* are predicted to be pathogenic. Structured domains are most sensitive (Figure S2A), while the most common benign mutations occur in IDRs (Figure S2B). All variants currently associated with congenital microcephaly exist within or lead to absence of conserved structured domains (Figure 1A). We chose six variants to profile (A109P, G201W, V229G, R236X, R536C, and Q782X), based on a combination of clinical information, site conservation, and previous *in vivo* experimentation validating LOF in brain development ^5,6^ (Figure 2A). We used the same AP-MS approach as before. Expression of variants and their pulldown efficiencies were comparable between each condition (Figure S1B-C). Raw ANKLE2 protein quantities were also comparable between variants, except for R236X, likely due to fewer tryptic peptides (Figure S1D). Proteomic scoring and HC-PPI filtering for our six variants were performed using the same parameters as WT ANKLE2 (Figure 2B). Notably, the decrease in R236X abundance did not correlate with a decrease in identified interactions. Instead R236X had far more HC-PPIs than any other variant.

**Figure 2:**
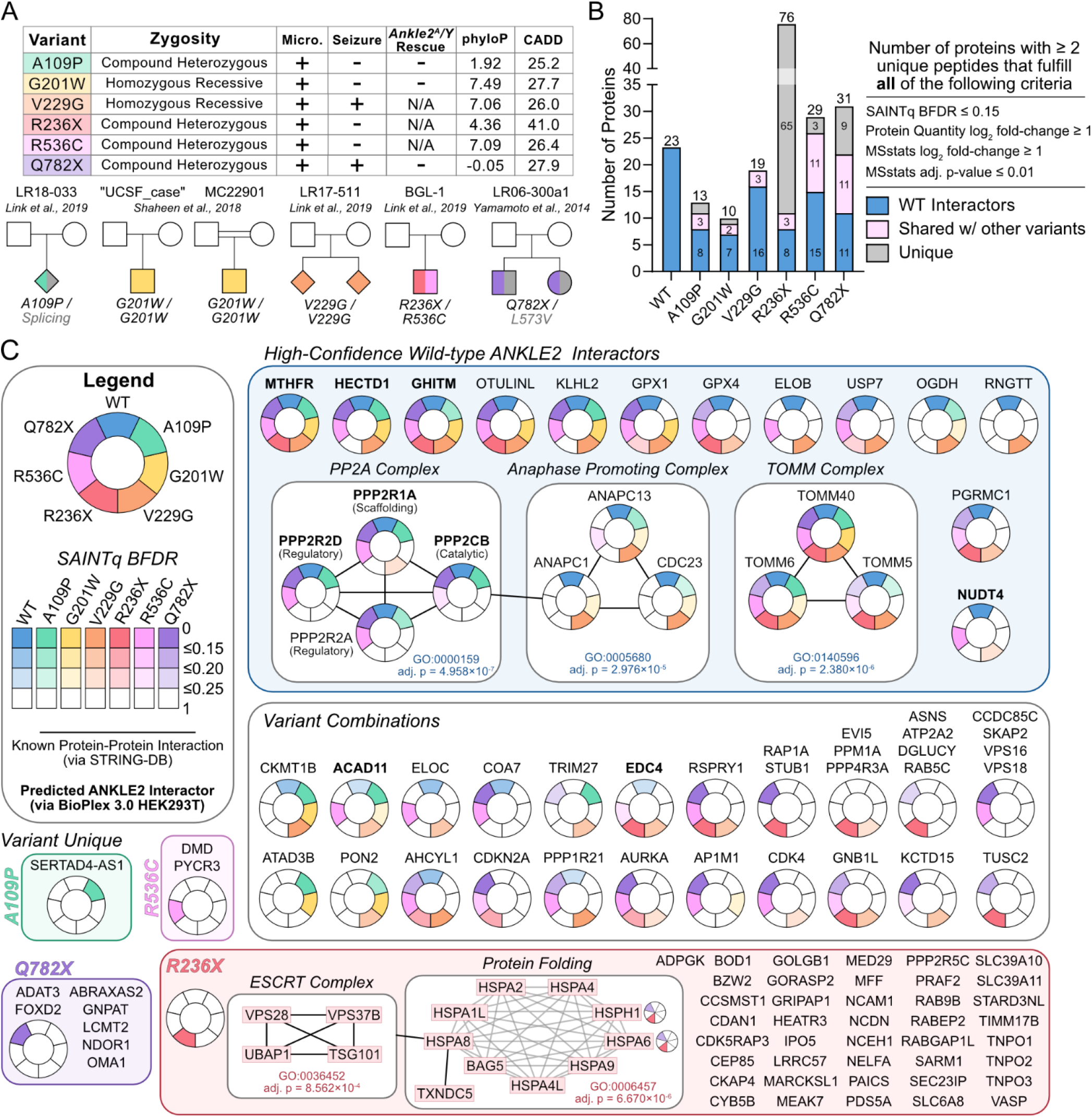
ANKLE2 pathogenic variants have altered protein interaction landscapes. **A)** Table and pedigrees of chosen pathogenic variants for evaluation in this study, based on allele, available clinical information ^4–6,13^. Micro: presence of clinical microcephaly; Seizure: clinical presence of seizures; *Ankle2^A^/Y* rescue: variant’s ability to rescue brain development phenotypes *in vivo*; phyloP: nucleotide level evolutionary conservation score (phyloP100way); CADD: Combined Annotation Dependent Depletion Score ^102^. Pedigrees: square = biological males, circles = biological females, diamonds = not disclosed, two horizontal lines = consanguineous couple, grey alleles = not evaluated in this study. **B)** Summary bar plot of ANKLE2 protein interactions for WT and pathogenic variants. Interacting prey proteins were evaluated using four criteria to determine high-confidence interactors (Figure S1). **C)** Visualization of ANKLE2 protein interactions for WT and pathogenic variants. Known protein interactions from STRINGdb are highlighted with black edge. Known ANKLE2 interactors identified by BioPlex are highlighted with a bold name. Confidence of interaction is indicated by color intensity and corresponds to SAINTq BFDR.

Using g:GOSt enrichment analysis on HC-PPIs, we observed clear differences in enrichment patterns between WT ANKLE2 and each pathogenic variant, especially for cell division processes (Figure S1G, Supplementary Data 1). V229G, was most similar to WT in its enrichment profile. The large truncation variant R236X had the most distinct profile, with unique enrichment in categories related to protein folding. Altogether, this analysis revealed a range of subtle to dramatic changes in the scaffolding function between WT ANKLE2 and variants associated with primary microcephaly.

We next visualized changes in the ANKLE2 interactome associated with each pathogenic variant (Figure 2C). To highlight that protein interactions may not be completely lost by different pathogenic variants, we used SAINTq BFDR (Bayesian False Discovery Rate) values as a proxy for interaction confidence, with low BFDR values indicating high confidence and high BFDRs indicating low confidence. All 23 WT HC-PPIs were identified with at least one variant, whereas each variant lacked multiple HC-PPIs found with WT. We identify some protein interactions seemingly unaffected by the mutations we tested, such as MTHFR (methylenetetrahydrofolate reductase) and GHITM (growth hormone inducible transmembrane protein), both of which are present in the BioPlex database of known ANKLE2 interactors.

We validated some of these interactions using co-IP (co-immunoprecipitation) and western blotting. We generally found co-IP results closely mirrored the general protein quantity values from our AP-MS (Figure S2C-J). For example, PP2A subunits showed decreased co-IP for variants G201W, V229G, and R236X, concordant with MS protein abundance data (Figure S2C-F). Only R236X failed to co-IP with HECTD1 (HECT domain E3 ubiquitin protein ligase 1), matching our AP-MS data where it was uniquely lost (Figure 2SC and H). TRIM27 co-IPed more strongly with A109P (Figure S2C and I). Finally, OMA1 (OMA1 zinc metallopeptidase) interacted strongest with Q782X as our proteomic data suggested (Figure 2SC and J). We did not identify any HC-PPIs unique to WT ANKLE2 and lost in all six variants, which would explain a universal mechanism for congenital microcephaly. Thus, we conclude that the ANKLE2 variants tested here disrupt different interactions yet result in the same disease outcome.

Several interactors displayed domain-specific trends, in which variants affecting the same structured domains lost similar interactions. We thus tested whether combining this information with eight N- and C-terminal ANKLE2 truncations could be used to definitively map the ANKLE2 domain mediating the interaction. The variants G201W and V229G are both in the CD and have similar loss of PP2A. Previous studies have shown interaction with PP2A relies on the CD (residues 197-252) and ANK (299-473) domains ^2,18^. In agreement with this, we found that interactions with PPP2R1A and PPP2CB were lost with truncation mutants lacking these domains, but not with mutants missing TM or LEM (Figure S2K-L).

WT ANKLE2 and the V229G variant had the most confident data regarding interaction with APC, although the other missense variants were close to our chosen cutoffs (Figure 2C). The clear difference in APC subunit protein quantities for Q782X suggests this interaction may be dependent on the structured region (Unc2) lost in this variant (Figure S2M-O, Supplemental Data 1). Unfortunately, we were unable to validate these trends or map the interaction domain by co-IP and western blotting due to lack of appropriate antibodies for APC.

We next evaluated ANKLE2’s interaction with the E3-ubiqutin ligase HECTD1. Using ANKLE2 truncations we showed that this interaction is mediated by the C-terminal half of ANKLE2, as it is lost in a Δ475-938 truncation, which lack Unc1 and Unc2. However, truncation Δ2-474, which contains both Unc1 and Unc2, and truncation Δ812-938, which contains Unc1 but not Unc2, still interact with HECTD1 (Figure S2K-L). Thus, we conclude that Unc1 is required for the interaction with HECTD1.

Our WT ANKLE2 interactome revealed a novel interaction with the membrane-resident TOMM complex (Figure 1C-D). This interaction was maintained in all variants analyzed (Figure 2C, S2C, S2G), none of which affect the TM domain of ANKLE2. We thus speculated that this interaction may be mediated by the ANKLE2 TM domain. N-terminal truncations lacking TM failed to pull down TOMM40, while all C-terminal truncations that have TM retained the interaction (Figure S2K-L). From this, we conclude that the ANKLE2 TM domain mediates an interaction with TOMM. Taken together, we identify ANKLE2 protein interactions that are lost or gained by pathogenic variants, and specific structured domains mediate these interactions (CD with PP2A, TM with TOMM, and Unc1 with HECTD1).

### GaMD simulations of ANKLE2 reveal strong intradomain correlations

We next sought to understand how ANKLE2 pathogenic variants drive global or local structural changes, which consequently impact their protein interactions. However, there are no experimentally resolved structures of ANKLE2. Its size and many unstructured regions make ANKLE2 a poor candidate for various experimental structural biology approaches. We thus took advantage of AlphaFold and GaMD simulations to predict the WT ANKLE2 protein structure and evaluate the dynamics of its structured domains before studying how mutations alter them ^38,39^.

We simulated full length WT ANKLE2 using GaMD to increase conformational sampling compared to conventional MD ^38^. We started by predicting the three-dimensional structure using ColabFold ^40^. However, the TM of this structure was occluded and incompatible with membrane insertion (Figure S3A). We therefore predicted the structure of ANKLE2 with the TM deleted (Δ1-34) and simulated it to relax the soluble domains (Figure S3B). Following initial relaxation, the TM domain was appended to the structure, and the resulting protein was embedded in a heterogeneous bilayer mimicking the ER membrane composition (55% POPC, 35% POPE and 10% Cholesterol) ^41^. After system equilibration, the protein-membrane complex was simulated in triplicate using GaMD for ∼0.5 microseconds (µs) per replica (∼1.5 µs total) to characterize the baseline dynamics of WT ANKLE2 (Figure 3A). Root-mean-square fluctuation (RMSF) analysis was performed to assess the flexibility of the structured domains across replicas (Figure 3B). The N-terminal domains (TM, LEM, and CD) of WT ANKLE2 exhibited the highest fluctuation, exceeding even the unstructured regions. In contrast, the uncharacterized domains (Unc1 and Unc2) displayed the lowest fluctuation.

**Figure 3:**
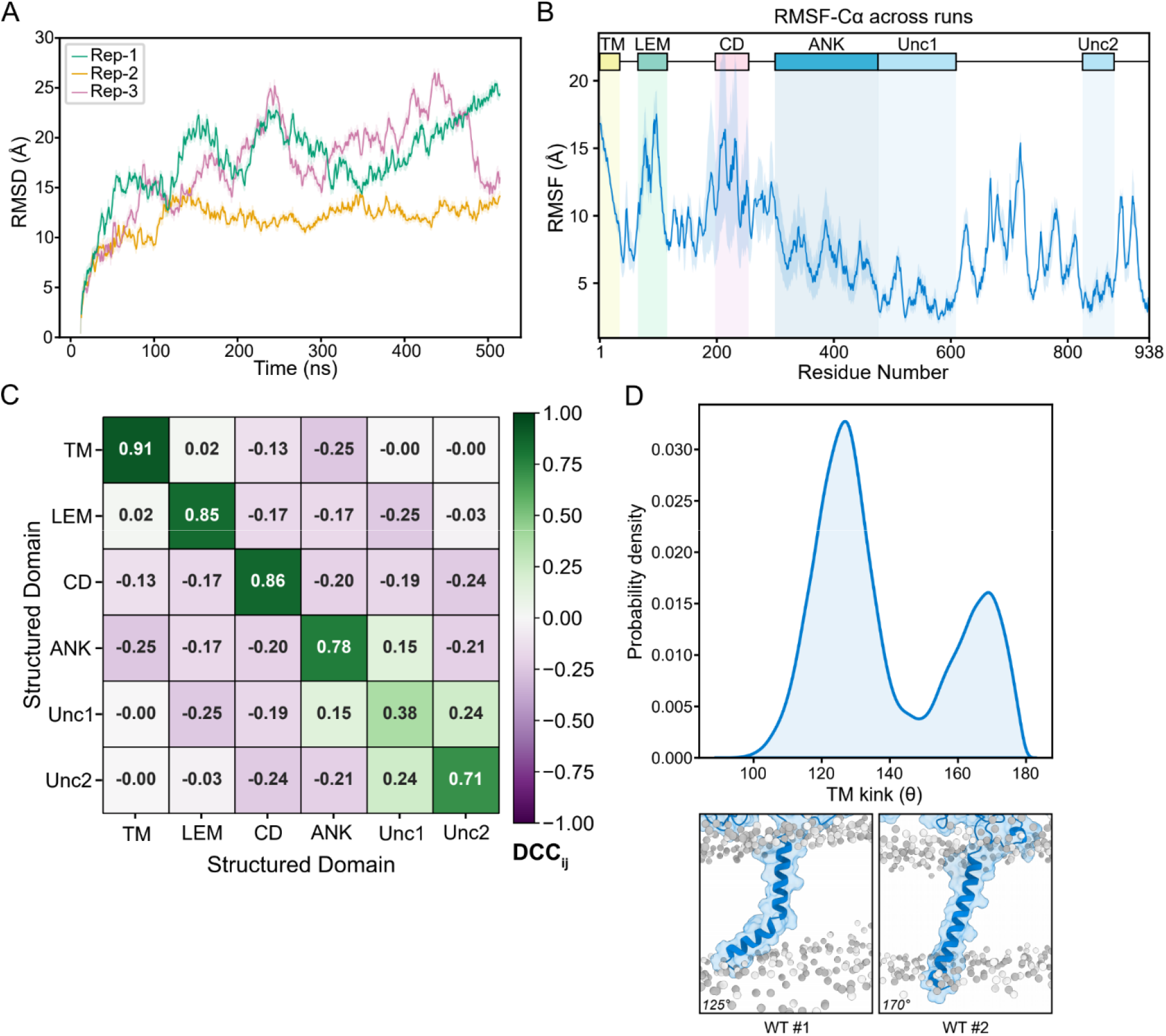
GaMD simulations of ANKLE2 reveals intradomain correlated motion and a TM kink. **A)** Root-mean-square deviation (RMSD) of ANKLE2 Cα atoms relative to the equilibrated structure across three independent replicas, colored by replica (Rep-1, Rep-2 and Rep-3). Muted lines show raw RMSD values at each frame, solid lines show the rolling average (window = 1 ns). **B)** Per-residue root-mean-square fluctuation (RMSF) of Cα atoms for ANKLE2 across three independent replicas. Solid lines indicate the mean RMSF, shaded regions indicate the standard deviation and residues corresponding to structured domains are annotated. **C)** Domain-averaged dynamic cross-correlation (DCC_ij_) among ANKLE2 structured domains (TM, LEM, CD, ANK, Unc1, Unc2), calculated for Cα atoms and block averaged to yield domain-specific correlation values (see Methods for details). **D)** Probability density of TM kink for ANKLE2 (top). Representative snapshots of TM domain within the bilayer illustrating the two dominant kink angles (∼125° and ∼170°) (bottom).

We next calculated dynamic cross-correlation (DCC_ij_) to assess how the structured domains of WT ANKLE2 influence one another’s motion. Positive values indicate correlated motions and negative values indicate anticorrelated motions between two Cα atoms i and j (Figure 3C, S3C) ^42^. Almost all structured domains showed strong intradomain correlation (>0.7). The only exception was Unc1 (+0.38), which alone showed correlated motion with the ANK (+0.15) and Unc2 (+0.24) domains. This suggests that Unc1 dynamics are weakly coupled with, or influenced by, motions in the ANK and Unc2 domains, rather than behaving as a fully independent structural unit. The motion of TM domain was largely decoupled from other domains, except for a weak anticorrelation with CD (−0.13) and ANK (−0.25).

Previously, we predicted the TM domain of ANKLE2 using DeepTMHMM (residues 10-32) ^15,43^, which modeled 23 residues of the TM domain (residues 1-34) as embedded within the bilayer. To further characterize these predictions, we examined the dynamics and orientation of the TM domain within the bilayer over the course of the simulations. TM kink analysis revealed that the helix predominantly adopts a kink angle of ∼125°, reflecting a bend in the helix, with a secondary population at ∼170°, reflecting a near linear conformation (Figure 3D). Taken together, these simulations allow us to characterize both local and global structural features of WT ANKLE2.

### R236X results in altered correlated motion and TM kink

Having established WT ANKLE2 dynamics, we next asked whether mutations within structured domains drive detectable changes in protein dynamics (Figure 4A). We first performed protein-level simulations of the truncation mutant R236X for ∼1.5 µs across three independent replicas (∼0.5 µs each) (Figure S4A). Our proteomics data showed that R236X had the most distinct protein interaction profile compared to WT ANKLE2, including interactions with multiple heat shock proteins (Figure 2C) ^44^. To test whether R236X shows altered conformational dynamics relative to WT ANKLE2 (residues 1-235), we performed principal component analysis (PCA) and observed clear separation along PC1, the direction that maximizes variance of the projected data (Figure 4B). Per-residue RMSF analysis showed R236X fluctuations were comparable to, or even lower than WT ANKLE2 (Figure S4B). Solvent accessible surface area (SASA) analysis revealed higher overall SASA for R236X compared to WT ANKLE2 (residues 1-235, Figure S4C). Thus, our data support a model in which normally buried residues may be extended and exposed in R236X, promoting its interactions with heat shock proteins ^45^ (Figure S4C).

**Figure 4:**
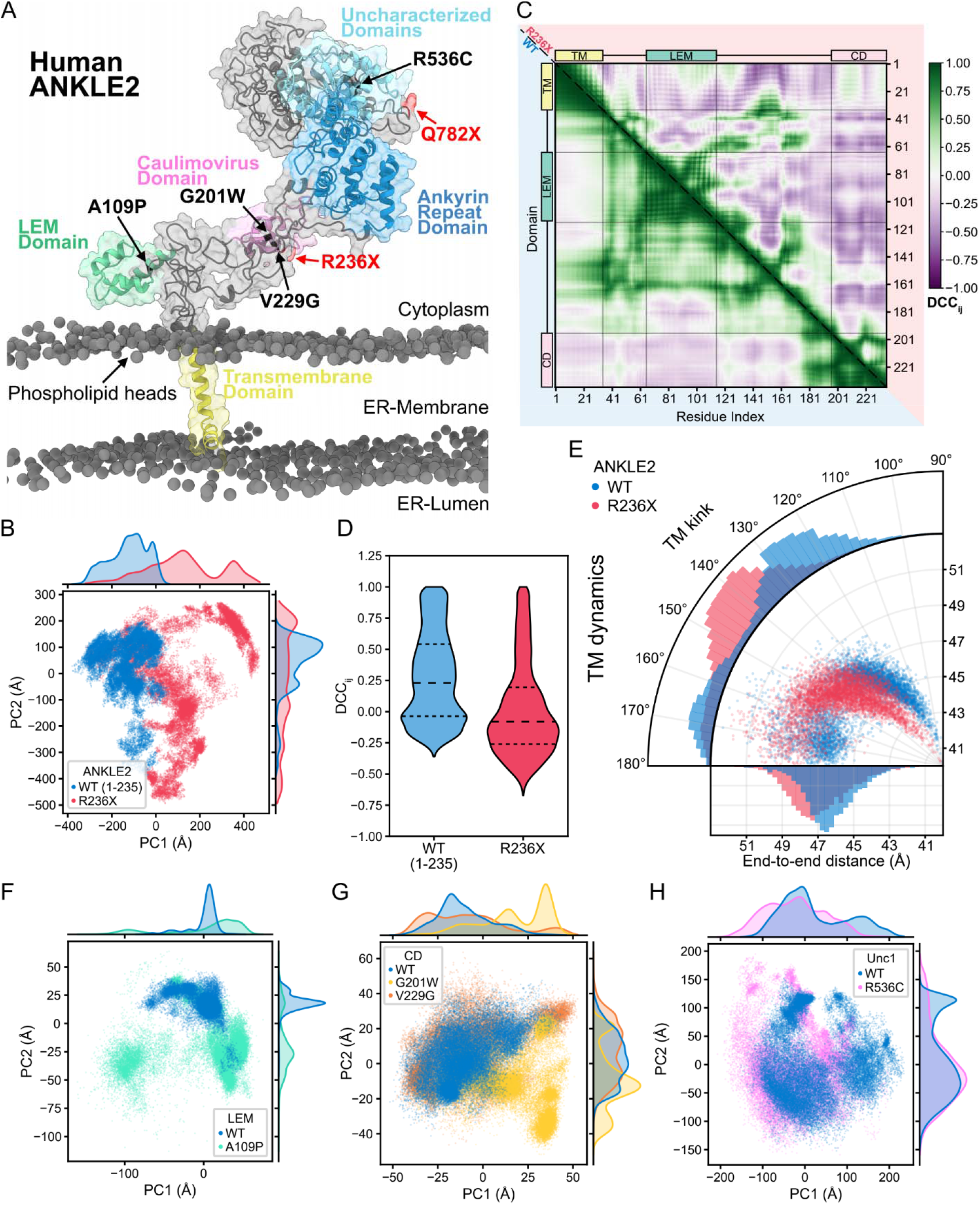
Altered intradomain communication and TM dynamics in nonsense mutant R236X. **A)** Structural model of ANKLE2 (Human ANKLE2) embedded in the membrane, showing phospholipid heads, annotated structured domains, and missense (black) and truncation (red) mutation sites. **B)** Principal component analysis (PCA) of ANKLE2 (residues 1-235) and R236X backbone atoms, across all replicas. **C)** Dynamic cross-correlation (DCC_ij_) for Cα atoms of ANKLE2 (residues 1-235) (lower matrix) and R236X (upper matrix), calculated across all replicas. The replica-averaged DCC_ij_ is shown. Residues corresponding to structured domains are annotated. **D)** Residues 1-235 Cα DCC_ij_ pooled across all replicas per mutant and shown as a violin plot (dashed lines represent quartiles). **E)** TM domain dynamics explained using TM kink and end-to-end distance pooled across all replicas per mutant and visualized as a radar plot. **F)** PCA of LEM domain WT and A109P backbone atoms backbone atoms, **G)** CD domain WT, G201W, and V229G backbone atoms, **H)** and Unc1 domain WT and R536C backbone atoms, across all replicas. Marginal kernel density estimates show the distribution of each mutant along PC1 (top) and PC2 (right).

Interestingly, comparison of DCC_ij_ values revealed that R236X loses global correlated motions compared to WT ANKLE2 (residues 1-235), and instead adopts predominantly anticorrelated motions (Figure 4C-D). This trend persisted within individual structured domains, with R236X showing reduced intradomain correlation relative to WT ANKLE2 (R236X – TM: +0.61, LEM: +0.58, CD*: +0.59) (Figure S4D). This reduced correlation was particularly notable given that the TM and LEM domains lie far from the truncation site. Because the TM domain was largely decoupled from other structured domains in WT ANKLE2 (Figure 3C), the reduced intradomain correlation in R236X warranted further investigation. Accordingly, we evaluated the TM dynamics in R236X using two structural metrics (TM kink and end-to-end distance) and found that R236X adopts a predominantly uniform TM kink of ∼145°, compared to the bimodal population of WT ANKLE2 TM at ∼125° and ∼170°. This shift was also reflected in the end-to-end distance of the helix, with R236X showing an increased distance relative to WT ANKLE2 (Figure 4E). Together, our dynamical analyses indicate that R236X has altered exposure of surfaces and long-range allostery, altering both its correlated motion and its behavior within the lipid bilayer. We propose that R236X is pathogenic because it is a large truncation whose dramatic structural changes disrupt many interactions, including those with HECTD1, PP2A, and APC.

To increase conformational sampling while reducing computational cost, we performed domain-level simulations of LEM (WT and A109P), CD (WT, G201W, and V229G), and Unc1 (WT and R536C) missense mutants. We simulated three independent replicas of 1 µs each (3 µs total). As a first step toward capturing high-level dynamical differences between WT and mutants, we performed PCA on the protein backbone across replicas. Some mutants had dynamics that were clearly separate from WT dynamics, while others were more subtle. For the LEM domain, A109P was clearly separated from WT along both PCs (Figure 4F). In CD, G201W differed from WT along PC1 and, to a lesser extent, PC2. In contrast, V229G showed less separation from WT, suggesting more subtle structural changes in CD for this variant (Figure 4G). In the Unc1 domain, R536C also showed little separation from WT in PC1 or PC2, suggesting minor changes in Unc1 domain dynamics (Figure 4H). Nonetheless, we were able to identify local structural changes related to R536C. Notably, WT Unc1 residue R536 participates in salt bridges with E493 and E533 (Figure S4E). These salt bridges can no longer form in R536C and may destabilize Unc1. We focused on the LEM domain and CD simulations moving forward because mutations in these domains resulted in larger global changes.

### The A109P mutation destabilizes the LEM domain structure

Given the dramatic shift in LEM PCA, we prioritized the A109P mutant for deeper analysis of the GaMD simulations. The LEM domain is predicted to have a characteristic three-helix fold (residues 65-74 (α1), 77-87 (α2), 99-111 (α3), Figure 5A) and A109P resides in α3. Root-mean-square deviation (RMSD) analysis revealed A109P diverged more than WT from its initial structure (Figure S5A). Per-residue RMSF analysis showed that A109P also had increased flexibility relative to WT across the LEM domain. The greatest divergence occurred in α3 (Figure 5A-B), suggesting that the proline substitution in α3 acts as a helix-breaker ^46^. Analysis of the charged residue interactions in α3 further supported this destabilization, as a salt bridge between R99 and E103 was stably maintained across the WT simulations but was notably less persistent in the A109P simulations, pointing to a loss of structural integrity in the mutant (Figure 5C).

**Figure 5:**
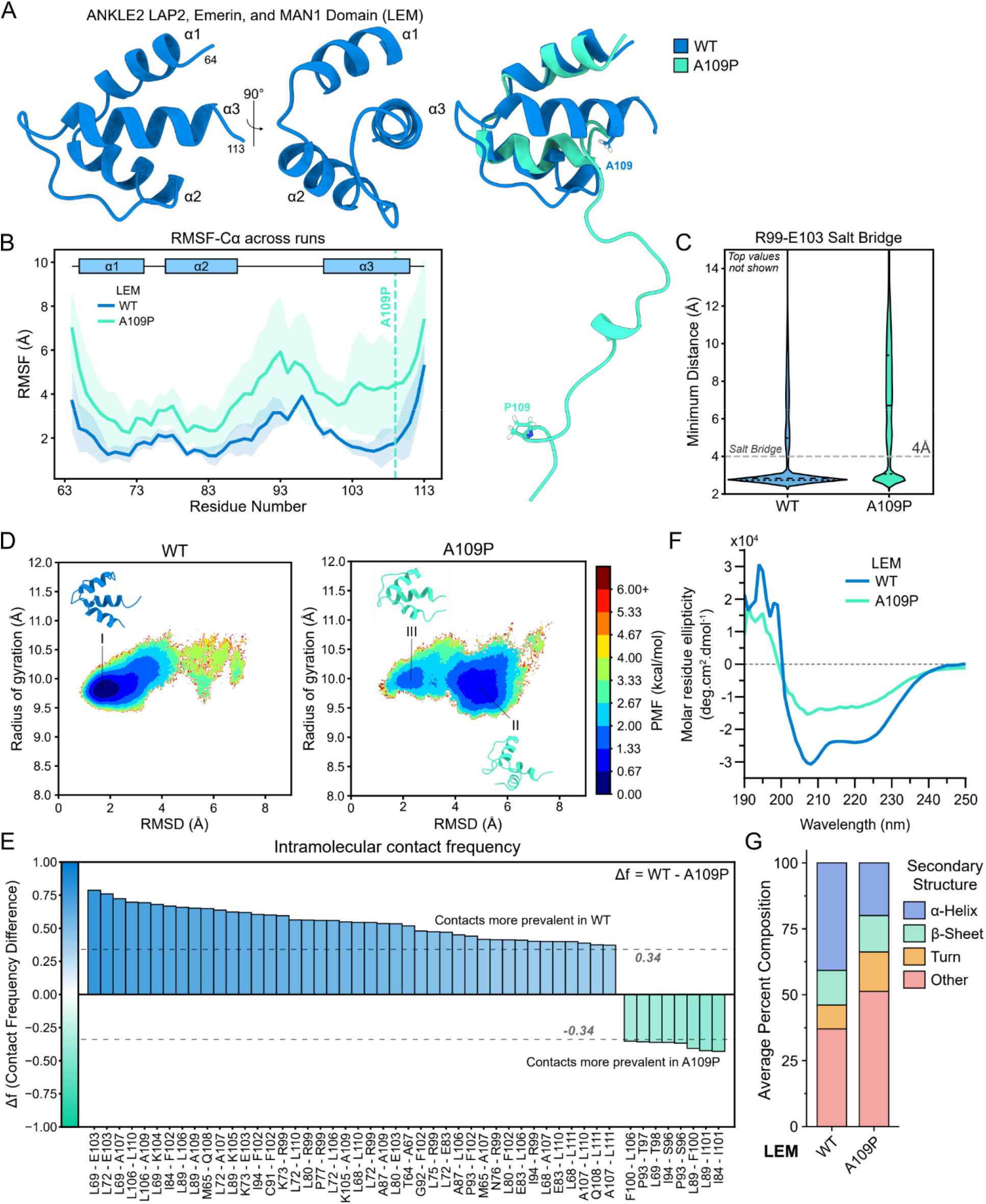
Proline substitution disrupts helical structure in the LEM domain. **A)** Ribbon representation of the ColabFold-predicted WT LEM structure, with residue numbers denoting domain boundaries and helices annotated (left), structural alignment of WT (blue) and A109P (green) frames extracted from the GaMD simulations (right). **B)** Root-mean-square fluctuation (RMSF) of Cα atoms for WT and A109P LEM domain across three independent replicas. Solid lines indicate the mean RMSF, shaded regions indicate the standard deviation. Residues corresponding to secondary structural elements and mutation site are annotated. **C)** Minimum distance between the nitrogen atoms of R99 and oxygen atoms of E103, pooled across all replicas per mutant and shown as a violin plot (dashed lines represent quartiles). The salt bridge distance cutoff (4 Å) is indicated. **D)** 2D potential of mean force (PMF) landscapes for WT (left) and A109P (right) LEM domain, reweighted individually per replica and aggregated, with root-mean-square deviation (RMSD) and radius of gyration (RoG) as the reaction coordinates. Representative structures from low-energy conformations shown for WT LEM (I) and A109P (II, III). **E)** Difference in reweighted binary intramolecular contact frequency (Δf) between WT and A109P LEM domain, pooled across replicas and shown as a waterfall plot for contact pairs with |Δf| > 0.34. Raw GaMD contact frequencies were reweighted using 10^th^ order Maclaurin series. **F)** Circular dichroism spectra of WT and A109P LEM domain following buffer subtraction plotted as molar residue ellipticity (deg.cm^2^.dmol^-1^). Spectra represents one of two replicates (Supplementary Data 2). **G)** Mean deconvoluted circular dichroism spectra (BeStSel) of WT and A109P LEM domain, with percent secondary structure composition ^101^.

Using RMSD and radius of gyration (RoG) as reaction coordinates, we reweighted the GaMD simulations to identify the preferred, low-energy conformational states of WT and A109P LEM structures ^47^. The resulting 2D reweighted potential of mean force (PMF) revealed WT adopts a primary low-energy state centered at 1.7 Å RMSD and 9.8 Å RoG (Figure 5D left, S5B top; conformation I, 0.017 kcal/mol). A109P showed a right-shifted minimum at 4.7 Å RMSD and 10.0 Å RoG (Figure 5D right, S5B bottom; conformation II, 0.712 kcal/mol), consistent with a partially disordered LEM domain conformation, along with a metastable state at 2.1 Å RMSD and 10.0 Å RoG (Figure 5D right, S5B bottom; conformation III, 1.489 kcal/mol) that closely resembles the low-energy conformation of WT LEM (Figure 5D).

To pinpoint the intramolecular interactions underlying this loss of structure, we analyzed binary contact and hydrogen bond (H-bond) patterns across the WT and A109P simulations. Binary contacts were defined as any interatomic distance < 4 Å between two residues, irrespective of the number of atom pairs involved, to avoid overweighting larger side chains. Reweighted contact frequency for each residue pair was calculated and summed across all replicas. A reweighted contact frequency difference (Δf) > 0.34 indicates that the change in residue contact between WT and A109P was observed in more than one replica. This analysis identified 50 reproducible contact changes, with A109P losing 42 native contacts and gaining 8 mutant-specific contacts (Figure 5E). Strikingly, 44 of 50 contact pairs involved at least one residue in α3, indicating that the loss of helicity in the LEM domain is driven by structural rearrangement within this helix (Figure S5C). H-bond analysis provided further support for this, identifying 8 residue pairs with > 10% difference in reweighted H-bond frequency between WT and A109P. Six of these H-bonds included at least one residue in α3, with A109P showing reduced H-bond propensity at these positions (Figure S5D, Supplementary Video 1). Together, our simulations show that α3 structure is disrupted in A109P.

We sought to validate these computational findings experimentally. We recombinantly expressed and purified WT LEM and the A109P mutant to assess their secondary structure using circular dichroism spectroscopy. Both spectra displayed characteristic negative peaks at 208 nm and 222 nm indicative of α-helical structure. However, the A109P spectrum showed reduced amplitude relative to WT LEM, consistent with a decrease in helical content (Figure 5F-G).

Despite these large structural changes in LEM, the A109P variant still maintains interactions with PP2A and APC, unlike every other variant analyzed for protein interactions. This finding suggests that these changes in LEM structure do not propagate to larger changes in CD or other domains that would impact PP2A and APC interactions. Thus, structural changes caused by A109P likely affect other protein interactions pertinent to brain development.

### Structural characterization of CD mutants with impaired PP2A binding

We next considered how mutations in CD alter its dynamics. CD is a 55 residue domain is predicted to have two α-helices and three β-strands (ANKLE2 residues 199-203 (β1), 217-219 (β2), 222-231 (α1), 236-240 (β3), 243-250 (α2), Figure 6A). Our proteomic data show that CD variants G201W and V229G disrupt binding to the PP2A complex members (PPP2R1A, PPP2CB, PPP2R2A and PPP2R2D) (Figure 2, S2C-F), consistent with CD’s known role in mediating the PP2A interaction ^2,18^. We hypothesized that these mutations disrupt local structure required for the PP2A interaction. We analyzed our GaMD simulations of WT, G201W, and V229G CD in more detail. RMSD analysis did not reveal any clear mutant-specific behavior (Figure S6A). Per-residue RMSF analysis showed that mutants exhibit flexibility similar to WT (Figure S6B).

**Figure 6:**
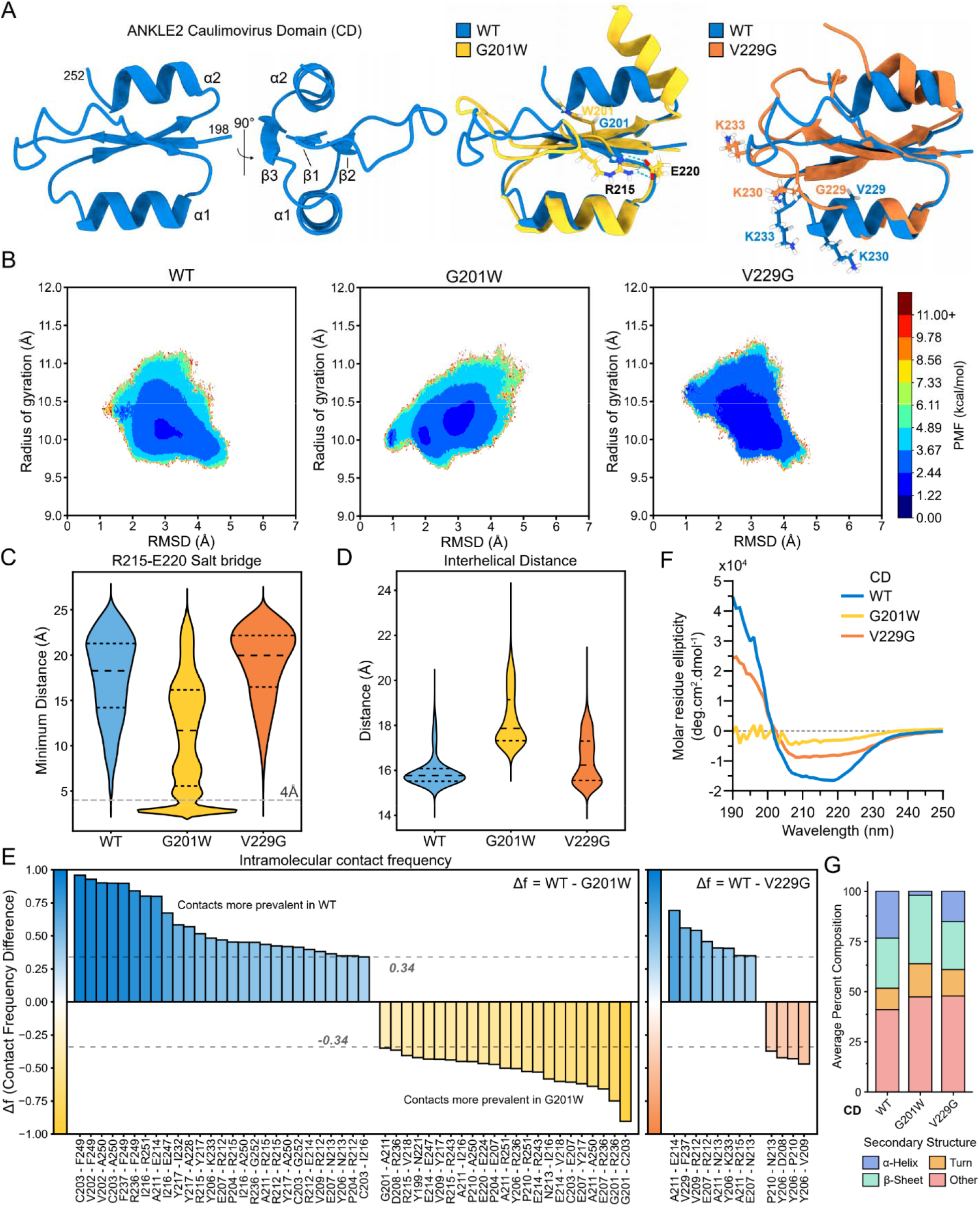
G201W and V229G induces widespread and subtle conformational changes in the CD domain, respectively. **A)** Ribbon representation of the ColabFold-predicted WT CD structure, with residue numbers denoting domain boundaries and secondary structures annotated (left), structural alignment of WT (blue) and G201W (yellow) or V229G (orange) CD frames extracted from the GaMD simulations (right). **B)** 2D potential of mean force (PMF) landscapes for WT (left), G201W (middle), and V229G (right) CD, reweighted individually per replica and aggregated, with root-mean-square deviation (RMSD) and radius of gyration (RoG) as the reaction coordinates. **C)** Minimum distance between the nitrogen atoms of R215 and oxygen atoms of E220, pooled across all replicas per mutant and shown as a violin plot (dashed lines represent quartiles). The salt bridge distance cutoff (4 Å) is indicated. **D)** Distance between the center of mass of the Cα atoms (representing the geometric center) of α1 and α2 within the CD domain, pooled across all replicas per variant. **E)** Difference in reweighted binary intramolecular contact frequency (Δf) between WT and G201W CD (left), WT and V229G CD (right), pooled across replicas and shown as a waterfall plot for contact pairs with |Δf| > 0.34. Raw GaMD contact frequencies were reweighted using 10^th^ order Maclaurin series. **F)** Circular dichroism spectra of WT, G201W, and V229G CD following buffer subtraction plotted as molar residue ellipticity (deg.cm^2^.dmol^-1^). Spectra represents one of two replicates (Supplementary Data 2). **G)** Mean deconvoluted circular dichroism spectra (BeStSel) of WT, G201W and V229G CD, with percent secondary structure composition.

To more precisely dissect these structural changes, we reweighted the GaMD simulations to identify the conformational states preferred by WT compared to mutants. The resulting 2D PMF landscape showed that WT had a minimum at 1.7 Å RMSD and 10.3 Å RoG (1.525 kcal/mol) and a metastable state at 2.9 Å RMSD and 10.1 Å RoG (2.225 kcal/mol) (Figure 6B, S6C, left). G201W had a minimum at 0.8 Å RMSD and 10 Å RoG (0.840 kcal/mol) along with two metastable states at 3.0 Å RMSD and 10.2 Å RoG (1.828 kcal/mol) and 1.9 Å RMSD and 10.0 Å RoG (2.108 kcal/mol) (Figure 6B, S6C, middle). V229G had a minimum at 1.2 Å RMSD and 10.7 Å RoG (1.031 kcal/mol) and a metastable state at 3.3 Å RMSD and 10.1 Å RoG (1.553 kcal/mol), along with a broad low-energy well spanning 2-4 Å RMSD and 9.8-10.7 Å RoG (Figure 6B, S6C, right). Together, these results indicate that, relative to WT, CD mutants access multiple discrete metastable states (G201W) and freely sample a wide range of conformations within a broad basin (V229G).

Given that the G201W mutation introduces a bulky side chain into β1, we examined its local structural effects in greater detail. Analysis of the charged residues within CD revealed that G201W mutant promotes formation of a new salt bridge between R215 and E220 (Figure 6C, S7A, Supplementary Video 2), potentially altering β-strand packing. G201W also alters the relative positioning of the two α-helices within the CD. V229G alters this same interhelical distance relative to WT, though to a lesser extent (Figure 6D, S7B).

We next analyzed binary contacts and H-bond patterns across WT, G201W, and V229G simulations to characterize changes in intramolecular interactions. Reweighted binary contact analysis identified 50 reproducible changes between G201W and WT, with G201W losing 27 native contacts while gaining 23 mutant-specific contacts (Figure 6E, left). Of these 50 contact pairs, 21 involved at least one residue in β1 or α2. Consistent with this, H-bond analysis identified 15 residue pairs with a > 10% difference in reweighted H-bond frequency between WT and G201W, 10 of which included at least one residue in β1 or α2 (Figure S7C). In contrast, binary contact analysis between V229G and WT yielded only 12 contact changes, including 4 mutant-specific contacts (Figure 6E, right).

As with the LEM domain mutant, we validated these observations experimentally. Recombinant WT, G201W, and V229G CD were purified and secondary structure was analyzed by circular dichroism spectroscopy. The G201W spectrum exhibited stark differences from WT across all wavelengths, translating to a loss of helical content relative to WT, likely driven by rearrangements within α2. V229G differed from WT in magnitude but largely maintained the overall spectral shape, consistent with partial structural perturbation of V229G (Figure 6F-G).

Collectively, data from proteomics, simulations, and experimental validation suggest G201W has more dramatic changes in protein interactions and structure than V229G. G201W exhibits loss or partial loss of interactions with PP2A and APC, along with drastic local structural rearrangement. In contrast, V229G is a more subtle mutation, inducing limited structural changes, that nonetheless results in loss of interaction with PP2A (Figure 2C). Together, these findings suggest that minor structural changes within CD result in pathogenic loss of interaction with the PP2A complex.

### ANKLE2 regulates the phosphoproteome through PP2A

ANKLE2 regulates BAF phosphorylation in a PP2A-dependent manner ^1,2,18,48^. Additionally, our ANKLE2 interactome revealed interactions with APC; a known cell cycle regulator, PP2A interaction partner, and putative microcephaly complex (Figure 1C). We therefore tested if ANKLE2 may broadly regulate phosphorylation of target substrates involved in brain development in a PP2A-dependent manner.

We identified ANKLE2-dependent and PP2A-dependent changes in protein phosphorylation using a phosphoproteomic approach (Supplementary Data 3). ANKLE2-dependent changes were identified comparing control and ANKLE2 KO cells ^31^. PP2A-dependent changes were identified using okadaic acid (OA), a potent inhibitor of all PP2A activity ^49^ (Figure 7A, S8A). As validation of our system, we measured phosphorylation of GEFH1, a known target of PP2A ^50^. We observed increased GEFH1 phosphorylation upon the addition of OA (Figure 7B). Neither ANKLE2 KO nor treatment with OA directly altered PP2A or BAF abundance (Figure 7B). OA also altered several phosphorylation sites on ANKLE2 itself (Supplementary Data 3). Treatment with OA in control cells revealed robust differences in global phosphorylation, whereas ANKLE2 KO resulted in a smaller set of significant changes, as expected (Figure 7C, S8B).

**Figure 7:**
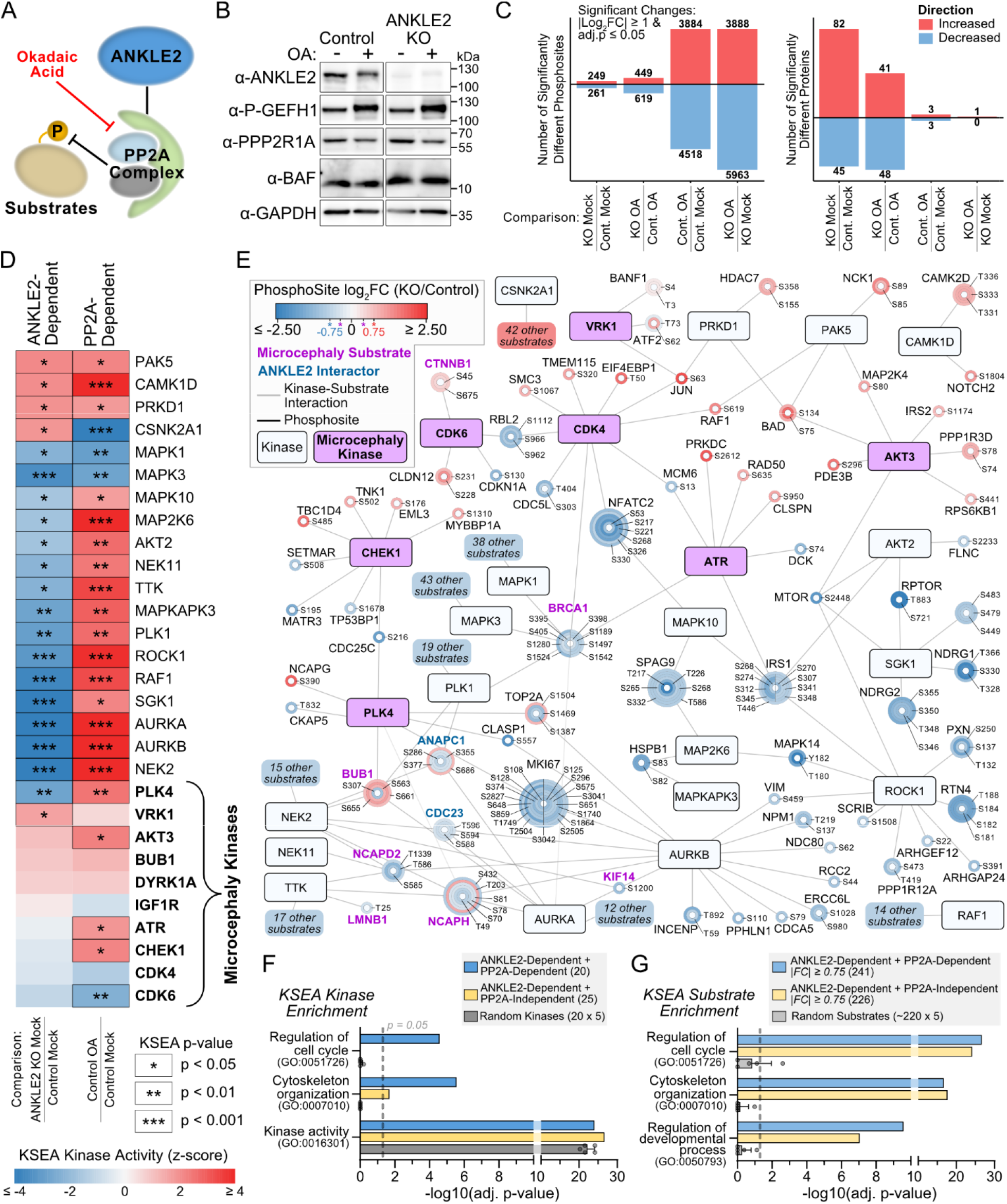
ANKLE2- and PP2A-dependent regulation of the phosphoproteome. **A)** Model of ANKLE2 interacting with the PP2A complex that regulates the phosphorylation state of cellular substrates. Okadaic acid (OA) is used to globally inhibit all PP2A complexes, allowing differentiation between effects dependent on ANKLE2 through PP2A and those independent of PP2A. **B)** Western blot of control and ANKLE2 KO cells treated with OA or DMSO. GEFH1 is a known target of PP2A and is used as a control to validate OA treatment. ANKLE2 KO does not inherently alter PP2A abundance. **C)** Counts of phosphosites (left) and total protein (right) that are considered statistically different between conditions using |log_2_ fold change| ≥ 1 and adjusted p-value ≤ 0.05 as criteria. **D)** Kinase-substrate enrichment analysis (KSEA) ^51^ was used to identify kinases with significantly altered activity dependent on ANKLE2. Known microcephaly genes (bold) from curated list (Supplemental table S1) are also included. **E)** Kinase-substrate network of ANKLE2-dependent and PP2A-dependent kinases with matching directional substrate |log_2_ fold changes| ≥ 0.75 or microcephaly kinases (pink boxes) with bidirectional |log_2_ fold changes| ≥ 0.75. Microcephaly substrates (pink) and identified ANKLE2-interactors (blue) with |log_2_ fold changes| ≥ 0.40 are also shown. Kinases associated with more than 200 identified phosphosites (m ≥ 200: CSNK2A1, MAPK3, NEK2, MAPK1, AURKA, TTK, RAF1, and PLK1) were limited to only microcephaly or ANKLE2-interacting substrates. The number of these hidden substrate proteins, irrespective of number of phosphosites or other kinase interactions, is indicated in colored boxes that correspond to kinase z-score. **F)** Functional enrichment of kinases from KSEA. ANKLE2-Dependent are all kinases with KSEA p-values < 0.05 for KO-Mock vs. Control-Mock. PP2A-Dependent are all kinases with KSEA p-values < 0.05 for Control-OA vs. Control-Mock. As a control for enrichment 20 kinases were randomly selected 5 times from the 395 kinases in the KSEA dataset. **G)** Functional enrichment of substrates from KSEA, based on the kinase targets from F). Substrate phosphosites with |log2 fold changes| < 0.75 were excluded. As a control for enrichment ∼220 substrates were randomly selected 5 times from the 7008 substrates in the KSEA dataset.

To dissect how ANKLE2 dysregulates cellular phosphorylation, we used kinase-substrate enrichment analysis (KSEA) to identify kinases with significantly altered behavior based on the profile of their known substrates ^51^. When we compared ANKLE2 KO cells and control cells we identified 29 kinases with significantly reduced enrichment and 16 kinases with significantly increased enrichment (Figure S8C). The kinases with increased enrichment in ANKLE2 KO cells corresponded to kinases inhibited by ANKLE2. Amongst these was VRK1, whose kinase activity on BAF is known to be inhibited by ANKLE2 ^2,5^. While BAF phosphorylation at T3 and S4 were detected and slightly increased in ANKLE2 KO cells, these changes were not significant. We performed motif-enrichment analysis ^52^ did not observe any consistent substrate motifs that may serve as preferential regulatory sites for ANKLE2 and interacting kinases. Based on the volume of other significantly different substrate phosphorylation events and kinases, we conclude that ANKLE2 may serve in scaffolding many kinase-substrate interactions.

We next analyzed the intersection of kinase-substrate interactions that were ANKLE2-dependent and PP2A-dependent. We identified 20 kinases that fulfilled these criteria (Figure 7D) and mapped their kinase-substrate pairings (Figure 7E). This set of kinases included PLK4, a key regulator of centriole biogenesis and microcephaly-associated gene ^53,54^. This prompted us to investigate if other kinases or substrates were associated with genetic disorders of microcephaly. To this end, we curated a list of 102 genes with clinical associations to primary microcephaly (Supplemental Table S1). We identified phosphorylation sites (phosphosites) associated with 55 microcephaly genes in our dataset and 10 kinases in our KSEA (Figure 7E, Supplementary Data 3). The phosphosites did not coincide with known pathogenic residues for these proteins ^55^. We also included the high-confidence ANKLE2-interactors ANAPC1 and CDC23 as putative microcephaly candidates from this study (Figure 1). Both ANAPC1 and CDC23 had multiple phosphosites with decreased signal in ANKLE2 KO cells (Figure 7E). During this analysis we also identified 25 kinases with significant ANKLE2-dependent enrichment that were not significantly enriched during OA treatment, suggesting these kinases may be regulated by ANKLE2 independently of PP2A (Figure S8C). In this set of ANKLE2-dependent and PP2A-independent kinase-substrate pairs, we identified 8 microcephaly genes, some of which we identified previously, but through different phosphosites (Figure S8D).

To gain insight into the functional processes controlled by ANKLE2-dependent phosphorylation, we performed additional functional enrichment on both the kinases and substrates we identified through KSEA (Supplemental Data 3). Background enrichment was calculated by randomly selecting a similar number of kinases or substrates from the database. Enrichment of ANKLE2-dependent and PP2A-dependent kinases revealed significant enrichment for regulation of cell cycle and other cell division processes, however this was far less apparent with PP2A-independent kinases (Figure 7F). When we evaluated the substrates, we observed significant PP2A-dependent and -independent enrichment of the same categories (Figure 7G). We also found enrichment for multiple developmental categories, supporting that ANKLE2 is a hub for the phosphoregulation of cell division and development (Figure 7G, Supplemental Data 3).

Taken together, our data proposes a role for ANKLE2 in regulating a wide range of phosphorylation events. By inhibiting PP2A with OA, we determine that many of these are mediated through ANKLE2’s interaction with the complex. We extrapolate from this that pathogenic variants that cannot maintain this interaction, such as G201W and V229G, will have similarly dysregulated phosphoproteomes. The breadth of proteins and phosphorylation sites impacted in this way points to a wide range of sources for ANKLE2-mediated congenital microcephaly.

## Discussion

The molecular and cellular consequences of pathogenic gene variants that ultimately result in congenital disease can be difficult to define mechanistically. This is especially challenging for non-enzymatic proteins lacking clear roles like ANKLE2. By integrating global proteomic approaches, molecular dynamics simulations, and validation *in vitro* and *in vivo,* we begin to unravel mechanisms for how specific variants cause neuropathogenesis.

Through our integrative approach, we provide a unifying framework for ANKLE2 pathogenic variants in this study. While there was no single interaction lost by all variants, interaction with either PP2A or APC was commonly lost except with A109P. We speculate the PP2A-APC axis is a critical hub for ANKLE2-dependent brain development that is commonly disrupted by pathogenic variants. PP2A orthologs in yeast can dephosphorylate APC and inhibit its function ^56^. Our data showed decreases in phosphorylation of APC subunits ANAPC1 and CDC23 (Figure 7E), suggesting APC phosphorylation state may be regulated in a PP2A- *and* ANKLE2-dependent manner. The functional outcome of phosphorylation at these sites is currently unknown, but could impact APC-mediated asymmetric cell division ^57^, a process also perturbed by *Ankle2* LOF mutants in flies ^5^. Moreover, while we suggest APC subunits as candidate microcephaly genes, it is important to note that the essential role of APC in neurogenesis and embryonic development is not new to our study ^58–62^. We propose that subtle and non-lethal APC mutations exist that dysregulate brain development to cause microcephaly, similar to ANKLE2 ^4–6^.

A109P did not disrupt interactions with PP2A or APC, despite the dramatic local structural changes we observed. We speculate that, instead of acting through a PP2A-APC axis, A109P may be pathogenic because loss of interaction with BAF, which binds to the LEM domain and is critical for cell cycle progression ^1^. Experiments with improved BAF detection would offer mechanistic explanation for the A109P phenotype. Alternatively, other proteins that are uniquely lost by A109P could mediate its pathogenesis. OTULINL (OTULIN-like) is specifically lost by A109P. While OTULINL is not well characterized, the related OTULIN is a key regulator of inflammation and angiogenesis during development ^63^. We speculate OTULINL also plays important roles in development.

The union of proteomics with GaMD also provides nuanced insight into specific pathogenic variants. For example, aside from loss of interaction with PP2A, the pathogenic nature of V229G is surprising. The substitution itself replaces a small hydrophobic residue with another small hydrophobic residue. Additionally, its protein interaction profile and structural dynamics are very similar to that of WT (Figures 2 and 6). Nonetheless, even subtle changes in CD dynamics result in loss of interaction with PP2A. PP2A is crucial for brain function and development, and mutations in PP2A subunits cause neurodevelopmental disorders ^64,65^. Our integrative approach highlights the importance of PP2A as a central hub in ANKLE2 pathogenesis.

Our integrative approach also identified a previously unknown role of ANKLE2 in regulating mitochondrial function (Figure 1C-E). Mitochondrial dysfunction is associated with congenital diseases, including defects in embryonic neurogenesis ^66^. While we found mitochondrial membrane potential is regulated by ANKLE2, how this is connected to development is still unknown. Additionally, other emerging mitochondrial functions, including inter-organelle contacts and lipid trafficking ^67,68^ could be regulated by ANKLE2.

In addition to APC, our AP-MS study identified other ANKLE2 interactors that may be *bona fide* MCPH proteins themselves. HECTD1 is involved in brain development ^69^ and loss of *Hectd1* in mice also leads to microcephaly ^70^. While no human variants are associated with microcephaly, recent work has identified mutations that result in neural tube defects ^71^. Our findings show that Unc1, which has eluded functional assignment ^15^, is essential for interaction with HECTD1 (Figure S2K-L). We propose to rename Unc1 as the HECTD1-binding domain (HBD) based on this new functional information. We speculate that R236X and other known *ANKLE2* pathogenic variants that disrupt or exist within HBD (L573V, G566R, and G585V) may compromise this interaction. Though, how ANKLE2’s interaction with HECTD1 contributes to cellular or developmental pathways requires additional exploration.

Finally, our study suggests that the microcephaly protein universe is sparse but interconnected, and can be traversed best with a versatile toolkit that leverages complementary approaches. The only MCPH protein known to interact with ANKLE2 (Supplemental Table 1) was CDK4 ^72^. Yet, we identified ANKLE2 interactions with other proteins implicated in brain development and microcephaly, including MTHFR ^73^, APC, and HECTD1. We speculate that such links must be common. Similar protein interaction mapping studies with other MCPH proteins will provide an atlas to navigate brain development and how it is disrupted by rare gene mutations. Integrating them with atomistic simulations and *in vivo* models will accelerate these mechanistic discoveries.

### Limitations of the study

Despite the many complementary techniques leveraged in this study, limitations remain. Namely, our AP-MS data excelled in identifying high-confidence ANKLE2 interactions that we could reliably validate, but our study was performed in HEK293T cells. While many of the interactions were relevant to brain development, future efforts to map ANKLE2 interactions in neuroprogenitor cells could reveal additional biologically relevant interactions. Additionally, our AP-MS study failed to reproduce the canonical ANKLE2 interactors BAF and VRK1 with high confidence ^1,2^. Both proteins are detected in the HEK293T dataset, but have no enrichment compared to GFP controls (Supplemental Data 1). Lack of enrichment of BAF may be explained by its highly entangled nature in the nuclear lamina. BAF purification requires more harsh disruption, in the form of sonication and/or nuclease digestion ^74^, that we were unable to perform here. However, even in AP-MS studies on BAF which employ these approaches, they fail to identify ANKLE2 ^74,75^, suggesting the interaction between them may be transient and cell cycle-dependent. The inability to detect this interaction does not detract from the proteins we were able to identify, rather it implies ANKLE2 may have other interactions we do not yet have the capacity to find. Similarly, interaction with VRK1 may be cell cycle-dependent and require specialized experimental setups.

While recent advances in protein structure prediction have been substantial, AlphaFold2-based models are known to perform poorly on intrinsically disordered or flexible loop regions. An experimentally derived ANKLE2 structure would offer a more reliable starting point for contextualizing the dynamics observed in our full length ANKLE2 simulations. Additionally, we simulated isolated structured domains that provide local structural context for individual mutations. While this approach focuses our attention on regions with high confidence in predicted structure, it will not capture global conformational changes. This complicates direct comparison to our proteomic data, since loss or gain of interaction may be mediated by changes in multiple domains rather than by any single domain.

Finally, our phosphoproteomic studies use OA, which is a global inhibitor of PP2A and PP1. There are many PP2A complexes with different regulatory roles beyond the ANKLE2-interacting complex comprised of PPP2R1A, PPP2R2D, and PPP2CB. While only these subunits are proposed as high-confidence ANKLE2 interactors based on both our data and literature ^2,18^, we cannot exclude the possibility that ANKLE2 may indirectly regulate PP1 or other PP2A complexes. Additional experiments are required to explore the downstream consequences induced by this impaired phosphoregulation.

## Resource availability

### Lead Contact

Further information and requests for resources and reagents should be directed to and will be fulfilled by the lead contact, Priya S. Shah.

### Material availability

Plasmid constructs generated in this study are available from the lead contact with a completed material transfer agreement.

### Data and code availability

- Mass spectrometry data has been deposited in the ProteomeXchange Consortium and is accessible with the dataset identifiers PXD083005 and PXD084708.
- Materials related to molecular dynamics simulations are available from the lead contact upon request.
- Any other additional information required to reanalyze the data reported in this paper is available from the lead contact upon request.

## Acknowledgements

This work was supported by funding to P.S.S. provided by the University of California, Davis, and the National Institutes of Health (NIH) (R01AI170857). Funding was provided to J.R.J. by the NIH (R01AI167691). I.J.H. was supported by NIH (5R25NS130964). J.L.A.F. was supported by a National Ataxia Foundation fellowship and is a W. M. Keck Foundation Fellow. J.H.B. is supported by a W. M. Keck Foundation Grant and the NIH (R35GM163701). P.R.B. and S.A. acknowledge support from the ACS Petroleum Research Fund (68836-DNI6). A.P.G. and D.J.M. acknowledge support from the NIH (R35GM166109). We thank the UC Davis Proteomics Core staff, including Brett Phinney and Michelle Salemi for crucial work related to this project. We thank the UC Davis Protein Structure and Dynamics Core, including Dr. Madhu Budamagunta, for circular dichroism measurements. This work used Expanse at the San Diego Supercomputing Center through allocation BIO250052 from the Advanced Cyberinfrastructure Coordination Ecosystem: Services & Support (ACCESS) program, which is supported by U.S. National Science Foundation grants #2138259, #2138286, #2138307, #2137603, and #2138296 ^76^. We thank members of the Shah lab for their constant support and helpful feedback.

## Author contributions

A.T.F., V.P., and P.S.S. conceived the project and designed experiments. A.T.F., N.J.L., C.J.F., and M.L.R. prepared AP-MS samples. A.T.F., C.L.S., and P.S.S. analyzed AP-MS data. A.T.F., N.J.L., C.J.F., V.P., M.L.R., S.S.B., and B.L.G. performed validation experiments. V.P. performed molecular dynamics simulations with input from P.R.B., S.A., and P.S.S.. I.M.B. performed protein purification for circular dichroism spectroscopy with input from V.P., A.P.G., and D.J.M.. A.T.F., H.P., B.B., and J.R.J. performed and analyzed phosphoproteomics. I.J.H., E.M., and N.L. performed assays in flies. A.T.F., E.M., S.S.B., J.L.A.F., J.H.B., performed mitochondrial activity assays. A.T.F., V.P., P.R.B., N.L., and P.S.S. visualized data. P.S.S. supervised research and acquired funding. J.H.B., D.J.M., S.A., N.L., and J.R.J. provided additional resources and expertise. A.T.F., V.P., and P.S.S. wrote the manuscript.

## Methods

### Cells

HEK293T (gift of Dr. Sam Díaz-Muñoz), Huh7 (gift of Dr. Raul Andino), and Vero (ATCC) cell lines were maintained in Dulbecco’s modified Eagle’s medium (DMEM, Gibco ThermoFisher) supplemented with 10% fetal bovine serum (FBS, Gibco ThermoFisher) at 37°C, 5% CO_2_. Cells we washed with Dulbecco’s phosphate buffered saline (D-PBS, Life Technologies) and dissociated with 0.05% Trypsin-EDTA (Life Technologies). Cells were tested for *Mycoplasma* spp. monthly by PCR.

### Plasmids

ANKLE2 pathogenic variant DNA fragments were synthesized by Twist Biosciences and inserted into pcDNA4_TO, cut with KpnI and ApaI, with C-terminal 3xFLAG affinity-tags using Gibson assembly. Sequences for human LEM (WT and A109P), CD (WT, G201W and V229G) were codon optimized for expression in *E. coli.* Fragments were synthesized by Twist Biosciences and inserted into the pTB146-His6-SUMO expression vector cut with SapI and BamHI via Gibson assembly. Correct assembly was verified with whole plasmid sequencing (Plasmidsaurus).

### Transfection, affinity purification, and ANKLE2 interactome proteomic sample preparation

HEK293T cells were plated in 10 cm dishes and grown overnight (5×10^6^ cells in 10 mL media). Transfection was performed by combining 3.5 µg of each corresponding plasmid DNA with 700 µL of serum-free DMEM. Next, 21 µL of PolyJet transfection reagent (SignaGen) was combined with 700 µL serum-free DMEM and added to each plasmid DNA tube. Samples were mixed and incubated at room temperature for 15 minutes prior to addition to cells. Cells were then grown for an additional 24 hours. Transfection efficiency was confirmed using a GFP encoding plasmid. Media was then removed from each plate. To dissociate cells, 5 mL of D-PBS supplemented with 10 mM EDTA was added and allowed to incubate for several minutes. Cells were resuspended in 5 mL of D-PBS and transferred to 15 mL conical tubes prior to centrifugation at 94 g, 4°C for 5 minutes (Eppendorf centrifuge 5810 R, Rotor S-4-104). Cell pellets were washed with 5 mL D-PBS and centrifugation was repeated. Supernatant was removed and pellets were then resuspended in 1 mL IP lysis buffer (50 mM Tris Base, 150 mM NaCl, 0.5 M EDTA, pH 7.4) with Pierce^TM^ protease inhibitor tablets (Thermo Scientific) supplemented with 0.5% NP-40 Substitute (Igepal^TM^ CA-630, Affymetrix). Cells were lysed for 30 minutes at 4°C, and lysate was then centrifugated at 845 g, 4°C, for 20 minutes (Eppendorf centrifuge 5424 R, Rotor FA-45-24-11). Soluble protein was transferred to a fresh tube and stored at –80°C until all four biological replicates were prepared. Lysates were thawed on ice and a portion of each (60-100 µL) was collected, normalized by BCA assay (Thermo Scientific), and saved for western blotting analysis. Remaining lysate was added to 100 µL of magnetic FLAG beads (Sigma) and incubated overnight at 4°C with gentle rotation. Beads were then washed 4 times with 1 mL IP buffer with 0.05% NP-40 and once with 1 mL IP buffer without NP-40. A portion of beads were separated for protein elution using 40 µL of 100 ng/mL 3x FLAG peptide (APExBIO) at 211 g for 1 hour at room temperature (Eppendorf ThermoMixerC). Eluate was then removed, resuspended in NuPAGE LDS sample buffer and bond-breaker TCEP (Thermo Scientific), and boiled for 10 minutes at 95°C prior to evaluation by western blotting. The remaining beads were exchanged into 50 mM triethylammonium bicarbonate buffer (TEAB, Thermo Fisher). Proteins were digested on-bead in 500 ng of sequencing grade trypsin (Promega) overnight at 37°C, ∼1500 rpm. Digested peptides were collected and beads were incubated with an additional 200 ng trypsin for 2 hours at 37°C, ∼1500 rpm. Eluate peptides were combined with the previous collection prior to being stored at −80°C until further analysis.

### ANKLE2 interactome mass spectrometry and analysis

Digested peptides were directly loaded onto an Evosep C18 tip and separated using the Evosep One. Peptides were eluted and ionized using a Bruker Captive Spray emitter. A Bruker timsTOF Pro 2 mass spectrometer running in diaPASEF mode was used for acquisition. The acquisition scheme used for diaPASEF consisted of 6×3 50 m/z windows per PASEF scan. DIA data was searched using Spectronaut 17 (Biognosys) against the human (UP000005640, downloaded 5/10/2024) UniProt proteome along with a standardized contaminants database. The Direct DIA workflow was used under default settings. Briefly, trypsin/P Specific was set for the enzyme, allowing two missed cleavages and lengths between 7 to 52 amino acids. Fixed Modifications were set for Carbamidomethyl, and variable modifications were set to Acetyl (Protein N-term) and Oxidation. For DIA search identification, PSM and Protein Group FDR were set to 1%. A minimum of 2 peptides per protein group were required for quantification. Raw intensity values were normalized to all mapped peptides. Detected contaminants from non-human sources and ambiguous protein groups were removed.

Raw data was exported from Spectronaut (v20.4) and imported into RStudio (v4.3.1). A Spectronaut Protein Quantity Log_2_ fold change of ≥1 relative to negative controls (empty vector, GFP, and ER-GFP) was used as one criterion for determining a high-confidence interaction. MSstats (v4.10.1) ^25^ was used with standard settings including Log_2_ transformation, normalization set to equalize medians, and to remove proteins with one feature. MBimpute was set to ‘false’ and featureSubset was set to ‘all’. The groupComparison function was used to compare ANKLE2 or each variant to the combination of GFP and ER-GFP control samples. A MSstats log fold change of ≥1 and adjusted p-value of ≤0.01 were used as criteria for determining high-confidence interactions. SAINTq was used with standard settings in the fragment-level data mode with GFP and ER-GFP combined as the control group ^24^. A SAINTq BFDR cutoff of ≤ 0.15 was used in combination with previous criteria for determining high-confidence interactions. The mass spectrometry proteomics data have been deposited to the ProteomeXchange Consortium via the PRIDE ^77^ partner repository with the dataset identifier PXD083005.

### Phosphoproteomic sample preparation

ANKLE2 KO and control Huh7 cells (1e7) were plated in 15 cm dishes and grown to desired confluency. Cells were then treated with either 100 nM okadaic acid (ThermoFisher) or an equivalent volume of DMSO for 4 hours. Media was removed and cells were resuspended in D-PBS (10 mM EDTA) and pelleted as previously described. Cell pellets were resuspended in 1 mL of lysis buffer (8 M urea, 50 mM ammonium bicarbonate, 150 mM NaCl) and frozen at −80°C until four biological replicates were acquired.

Lysates were probe-sonicated on ice three times for 1 second at 50% power, with 5 seconds of rest in between pulses. Protein content of the lysates was quantified using a micro-BCA assay (Thermo Fisher Scientific). 1 mg of protein per sample was treated with Tris-(2-carboxyethyl)phosphine to a final concentration of 4 mM and incubated at room temperature (RT) for 30 minutes. Iodoacetamide (IAA) was added to each sample to a final concentration of 10 mM, and samples were incubated in the dark at RT for 30 minutes. IAA was quenched by dithiothreitol at a concentration of 10 mM and incubated in the dark at RT for 30 minutes. Samples were then diluted with five sample volumes of 100 mM ammonium bicarbonate. Trypsin Gold (Promega) was added at a 1:100 (enzyme:protein wt/wt) ratio and lysates were rotated for 16 hours at RT. 10% vol/vol trifluoroacetic acid (TFA) was added to each sample to a final concentration of 0.1% TFA. Samples were desalted under vacuum using Sep Pak tC18 cartridges (Waters). Each cartridge was first washed with 1 mL of 80% acetonitrile (ACN)/0.1% TFA, then washed three times with 1 mL of 0.1% TFA in water. Samples were then loaded on cartridges. Cartridges were washed three times with 1 mL of 0.1% TFA in water. Samples were then eluted with 1 mL of 40% ACN/0.1% TFA. 20 µg of each sample was kept for protein abundance measurements, and the remainder was used for phosphopeptide enrichment. Samples were dried by vacuum centrifugation. Protein abundance samples were resuspended in 0.1% formic acid (FA) for mass spectrometry analysis.

For each sample batch and under vacuum, 30 µL per sample of 50% Ni-NTA Superflow bead slurry (QIAGEN) was added to a 2-mL empty spin column (Bio-Spin, Bio-Rad). Beads were washed three times with 1 mL of HPLC-grade water, incubated four times with 1 mL of 100 mM EDTA for 30 seconds, washed three times with 1 mL of HPLC-grade water, incubated four times with 1 mL of 15 mM FeCl3 for 1 minute, washed three times with 1 mL of HPLC-grade water, and washed once with 1 mL of 0.5% vol/vol FA. Beads were resuspended in 750 µL of 80% ACN/0.1% TFA. 1 mg of digested peptides were resuspended in 83.33 µL of 40% ACN/0.1% TFA and 166.67 µL of 100% ACN/0.1% TFA, and 60 µL of the bead slurry was added to each sample and incubated for 30 minutes while rotating at RT. A C18 BioSPN column (Nest Group), centrifuged at 110 × g for 1 minute for each step, was equilibrated two times with 200 μL of 80% ACN/0.1% TFA. Beads were loaded on the column and washed four times with 200 μL of 80% ACN/0.1% TFA, then washed three times with 200 μL of 0.5% FA. Then, 200 μL of 500 mM potassium phosphate buffer pH 7 was added three times to the column and incubated for 1 minute. Then, 200 μL of 0.5% FA was added three times to the column. Phosphopeptides were eluted twice with 100 μL of 40% ACN/0.1% FA and vacuum centrifuged to dryness. Phosphopeptides were resuspended in 25 µL of 4% FA/3% ACN for mass spectrometry analysis.

### Phosphoproteomic mass spectrometry and analysis

These proteomic and phosphoproteomic samples were analyzed on an Orbitrap Eclipse mass spectrometry system equipped with an Easy nLC 1200 ultra-high pressure liquid chromatography system interfaced via a Nanospray Flex nanoelectrospray source (Thermo Fisher Scientific). Samples were injected onto a fritted fused silica capillary (30 cm × 75 μm inner diameter with a 15 μm tip, CoAnn Technologies) packed with ReprosilPur C18-AQ 1.9 μm particles (Dr. Maisch GmbH). Buffer A consisted of 0.1% Formic Acid (FA), and buffer B consisted of 0.1% FA/80% Acetonitrile (ACN). Analytical columns were equilibrated with 3 μL of buffer A. Peptides were separated by an organic gradient from 5% to 35% mobile buffer B over 120 min followed by an increase to 100% B over 10 min at a flow rate of 300 nL/min.

Phosphoproteome samples from each set of biological replicates were pooled and acquired in data-dependent acquisition (DDA) manner to build spectral libraries for data-independent acquisition data analysis. DDA data was acquired by collecting a full scan over a m/z range of 375-1025 in the Orbitrap at 120,000 resolving power (@ 200 m/z) with a normalized AGC target of 100%, an RF lens setting of 30%, and an instrument-controlled ion injection time. Dynamic exclusion was set to 30 seconds, with a 10 p.p.m. exclusion width setting. Peptides with charge states 2-6 were selected for MS/MS interrogation using higher energy collisional dissociation (HCD) with a normalized HCD collision energy of 28%, with 3 seconds of MS/MS scans per cycle.

Data-independent analysis (DIA) was performed on all individual samples. A full scan was collected at 60,000 resolving power over a scan range of 390-1010 m/z, an instrument controlled AGC target, an RF lens setting of 30%, and an instrument controlled maximum injection time, followed by DIA scans using 8 m/z isolation windows over 400-1000 m/z at a normalized HCD collision energy of 28%.

Proteome data were analyzed with a library-free approach using the DIA-NN algorithm ^78^. Data were searched against the Homo sapiens reference proteome sequences in the UniProt database (one protein sequence per gene, downloaded on August 23, 2023). Search parameters included a fixed modification for carbamidomethyl cysteine and variable modifications for N-terminal protein acetylation and methionine oxidation. All other search parameters were DIA-NN factory defaults.

For phosphoproteome data, the Spectronaut algorithm (version 15.2) was used to build spectral libraries from DDA data, identify peptides/proteins, localize phosphorylation sites, and extract intensity information from DIA data ^79^. All data were searched against the *Homo sapiens* reference proteome sequences in the UniProt database (one protein sequence per gene, downloaded on August 23, 2023). Data were filtered to achieve a false discovery rate of 0.01 for peptide-spectrum matches, peptide identifications, and protein identifications. Search parameters included a fixed modification for carbamidomethyl cysteine and variable modifications for N-terminal protein acetylation, methionine oxidation, and serine, threonine, and tyrosine phosphorylation. Peptide lengths from 7 to 52 amino acids were considered, and up to two missed cleavages were allowed. MS2 b and y ion types were utilized.

Statistical analysis of all proteome and phosphoproteome data was conducted utilizing the MSstats package in R ^80^. All data were normalized by equalizing median intensities, the summary method was Tukey’s median polish, and the maximum quantile for deciding censored missing values was 0.999, and multiple testing was corrected using the Benjamini-Hochberg procedure. For protein abundance analyses, only the top 50 features per protein were considered. This data was deposited to the ProteomeXchange Consortium via the PRIDE ^77^ partner repository with the dataset identifier PXD084708.

Phosphoproteomic data was analyzed using kinase-substrate enrichment analysis (KSEA) with the PhosphoSitePlus and NetworKIN (score cutoff = 2) datasets enabled ^51,81–83^. Kinases with p < 0.05 and m ≥ 3 in the ANKLE2 KO vs. Control DMSO conditions were then evaluated across all comparisons. Kinase-substrate networks were generated using Cytoscape (v3.10.4)^84^. For KSEA enriched kinases, only substrate phosphorylation sites with |log_2_ fold change| ≥ 0.75 in the same enrichment direction as the kinase were visualized. All phosphosites of microcephaly and ANKLE2-interactors with log_2_ fold change| ≥ 0.40 were visualized. Functional enrichment of these kinases and substrates were performed with gProfiler using the default g:SCS significance threshold ^28^. Background enrichment was controlled by randomly generating sets of kinases or substrates from the entire KSEA database.

### Western blotting and silver stain analysis

Protein samples (lysates or affinity-purification [AP] eluates) were resuspended in NuPAGE LDS sample buffer supplemented with tris(2-carboxyethyl)phosphine (TCEP) and boiled at 95°C for 10 min. Samples were run on 4%–20% gradient polyacrylamide gels for ∼1 hour at 150 V. For western blot proteins were transferred to polyvinylidene fluoride (PVDF) membranes (VWR) for 1 hour at 330 mA on ice. Membranes were then blocked in 5% milk solution for 1 hour prior to overnight incubation in primary antibodies (Table S1) at 4°C. Membranes were washed 3× in Tris-buffered saline with Tween 20 (TBS-T) (150 mM NaCl, 20 mM Tris base, 0.1% Tween 20; Thermo Fisher) and incubated with horseradish peroxidase (HRP) conjugated secondary antibodies in 5% milk for 1 hour at room temperature. Membranes were again washed 3× in TBS-T and 1× in Tris-buffered saline (without Tween 20) prior to Pierce ECL activation (Thermo Fisher). Membranes were imaged using Amersham Imager 600 (GE). For silver stain, the gel was prepared and stained per manufacturer’s recommendations (Thermo Fisher, PI24612). Gels were imaged with a BioRad GelDoc Go Imaging System. Images were analyzed using Fiji ^85^.

### Animal experiments

#### Fly lines and maintenance

The Drosophila melanogaster lines used in this study are the following: *D. melanogaster*: *P{VALIUM22-EGFP.shRNA.1}attP40*, RRID:BDSC_41557; *D. melanogaster*: *inscuteable-GAL4*: *P{w[+mW.hs]=GawB}insc[Mz1407],* RRID:BDSC_8751; *D. melanogaster*: *y[1] w[*]; P{w[+mC]=Act5C-GAL4}17bFO1/TM6B*, *Tb[1]*, RRID:BDSC_3954; *D. melanogaster*: *y[1] v[1]*; *P{y[+t7.7] v[+t1.8]=TRiP.HM04044}attP2*, RRID:BDSC_31735; *D. melanogaster*: *y[1] v[1]*; *P{y[+t7.7] v[+t1.8]=TRiP.HMC03337}attP40*, RRID:BDSC_51782; *D. melanogaster*: *y[1] sc[*] v[1] sev[21]*; *P{y[+t7.7] v[+t1.8]=TRiP.HMS01744}attP2/TM3*, *Sb[1]*, RRID:BDSC_38531; *D. melanogaster*: *y[1] v[1]*; *P{y[+t7.7] v[+t1.8]=TRiP.HMJ23608}attP40*, RRID:BDSC_61982; *D. melanogaster*: *y[1] w[*], P{ry[+t7.2]=neoFRT}19A*; *D. melanogaster*: y*[1] w[*] Ankle2[A] P{ry[+t7.2]=neoFRT}19A* / *FM7c*, *P{w[+mC]=GAL4-Kr.C}DC1*, *P{w[+mC]=UAS-GFP.S65T}DC5, sn[+]*, RRID:DGGR_117431; *D. melanogaster*: *y[1] w[*] Ankle2^CRIMIC^*. All stocks were raised at 25°C on Archon glucose formula food in wide vials.

#### *In vivo* RNAi

Females from ubiquitous knockdown driver line (*Actin5C-GAL4*) or neural stem cell knockdown driver line (*inscuteable-GAL4*) were crossed to males from *in vivo* RNAi lines targeted to APC or GFP for control. Crosses were set at 29°C on blue food which was made by adding bromophenol blue to the glucose food. Late third-instar larvae were selected for brain dissection based on extruding spiracles and gut clearance of blue food. *CG31687* included with *Cdc23* represents another likely homolog of CDC23. The inclusion of *fab1* with *Anapc13* is a limitation in the available fly lines.

#### Immunohistochemistry

Brains were dissected in phosphate-buffered saline (PBS) and transferred to microcentrifuge tubes for a 20-minute fixation with 4% paraformaldehyde in PBS + 0.3% Triton X-100 (PBST) and then washed with PBST three times for five minutes, twice with PBST + 5% Bovine serum albumin (PBSTB) for 30 minutes, and once with PBSTB + 5% Normal Donkey Serum for 30 minutes. Primary antibody (Deadpan, Abcam ab195173, neural stem cells) and pHH3 (phosphor-Histone H3, Millipore Sigma 06-570, mitotic cells) were incubated with samples overnight at 4°C. The following day, brains were washed three times with PBSTB for 20 minutes and incubated with secondary antibodies (Jackson ImmunoResearch, Donkey anti-rat Alexa fluor 647 Cat#712-605-153, RRID: AB_2340694 and Donkey anti-rabbit Alexa fluor 488 Cat#711-545-152, RRID: AB_2313584, 1:500) in PBSTB for one hour. Brains were washed three times with PBST and mounted in slow fade gold.

#### Brain lobe volume

A single brain lobe was imaged per brain on a Zeiss 710. The following confocal settings were used to image: 40X water immersion lens, zoom of 0.6, frame size of 1024 x 1024, z-slice size of 2 µm, and z limits were set using the Deadpan channel to capture the entire brain lobe. Using the surface function in Imaris, the perimeter of the brain lobe was traced using the Deadpan channel at every 5th z slice and then compiled to form a volumetric 3D surface. A multiple comparisons ANOVA was performed to identify statistical differences.

#### TMRM assay

Wandering third instar larvae were dissected (inverted) in filtered PBS and transferred to PBS with 200 nM TMRM for 10 minutes in the dark. Inverted larvae were briefly transferred to PBS and mounted in PBS. Samples were imaged within 1.5 hours on a Zeiss 980, and control was imaged first to set the base line. All confocal settings remained the same for each sample and each experiment except for z-stack. A variety of large muscle segments were imaged. One image per animal, 5-6 animals per experiment. Three independent experiments on different days. 20 µM FCCP was included with one experiment. Imaris was used to identify total sum of signal per z-stack and total volume of signal. Data is displayed as signal sum intensity divided by volume of signal.

### Molecular dynamics (MD) simulations

#### Modeling ANKLE2 structures

The initial ANKLE2 prediction was not compatible with membrane insertion of the TM domain. To generate a reliable initial relaxed structure of the soluble domains, we instead predicted ANKLE2 without the TM domain (ANKLE2 Δ1-34), along with individual ANKLE2 domains and mutants of these domains. ANKLE2 Δ1-34 (pLDDT=61.9), WT LEM (ANKLE2 residues 64-113, pLDDT=86.2), A109P (pLDDT=87.4), WT CD (ANKLE2 residues 198-252, pLDDT=84), G201W (pLDDT=81.6), V229G (pLDDT=83.3), WT Unc1 (ANKLE2 residues 472-620, pLDDT=87.4), and R536C (pLDDT=87.6) structures were modeled with ColabFold v1.5.5: Alphafold2 using MMseqs2 ^39,40,86^. Top five structures were relaxed using AMBER and top ranked structure was used for molecular dynamics (MD) and/or Gaussian accelerated molecular dynamics (GaMD) simulations ^38^.

#### ANKLE2 Δ1-34

ColabFold PDB files were processed and prepared using Schrödinger Maestro (release 2024-2). Missing hydrogen atoms were added, and protonation states were assigned at physiological pH 7.4 using protein preparation workflow ^87^. The processed ColabFold structures were parameterized using the AMBER ff19SB force field ^88^. Each protein structure was placed in a truncated octahedron box with a buffer of 20 Å and solvated using the OPC water model ^89^. The system was then neutralized and brought to physiological salt concentration (150 mM NaCl).

All the prepped systems were simulated using the following protocol. First, energy minimization was performed using the steepest descent algorithm for 1,000 cycles, followed by the conjugate gradient algorithm to relax the water molecules, counter ions and protein sidechains while keeping the protein backbone fixed with harmonic position restraints of 10 kcal.mol^-1^.Å^-2^. Next, a second energy minimization step was performed using the steepest descent algorithm for 1,000 cycles followed by conjugate gradient algorithm to relax the complete system. The systems were then heated up from 10 to 310 K in the canonical ensemble (NVT) over 5 ns and subsequently maintained at 310 K for an additional 5 ns, with position restraints of 1 kcal.mol^-1^.Å^-2^ imposed on the protein backbone.

Temperature control (310 K) was performed via Langevin dynamics ^90^, with a collision frequency of γ = 5.0 ps^-1^ during equilibration, later reduced to 1.0 ps^-1^ for production runs. Isotropic position scaling was applied for protein-only systems. Pressure was controlled by coupling the system to a Berendsen barostat ^91^, at a reference pressure of 0.987 atm with a relaxation time of 1 ps. All equilibration and production runs have been carried out using an integration time step of 2 fs and a non-bonded cutoff of 10 Å. Hydrogen atoms were added assuming standard bond lengths and were constrained to their equilibrium position with the SHAKE algorithm ^92^. All MD simulations were performed using pmemd.cuda module of AMBER 20 software suite ^93^. The final structures from the equilibration runs served as the starting point for the subsequent GaMD simulation.

#### WT ANKLE2 and R236X

Following initial relaxation of ANKLE2 Δ1-34, TM domain residues were generated using the start structure tool (Tools Structure Editing Build Structure Start Structure) and added to ANKLE2 Δ1-34 structure using join models tool in ChimeraX version: 1.6.1 ^94^ to perform ANKLE2 and R236X simulations. CHARMM-GUI ^95^ membrane builder was used to protonate the system at physiological pH 7.4, following which the transmembrane (TM) domain of ANKLE2 and R236X was embedded (the orientation of the protein was defined by two residues in the TM domain residue 6 and 27 along the z-axis) in a heterogeneous bilayer consisting of 1040 lipid molecules (55% POPC, 35% POPE and 10% Cholesterol) mimicking the composition of the endoplasmic reticulum ^41^. Protein-membrane complex was then solvated in explicit water molecules, 150 mM NaCl concentration and the system was neutralized with additional Na+ ions resulting in periodic simulation cells of ∼180*180*208 Å^3^ (815,512 atoms – ANKLE2) and ∼180*180*146 Å^3^ (559,624 atoms – R236X). CHARMM-GUI output systems were parameterized using the AMBER ff19SB, OPC and Lipid21 force fields. These protein-membrane systems were then used as input for GaMD simulations.

CHARMM-GUI prepped protein-membrane systems were simulated using the following protocol. First, energy minimization was performed using the steepest descent algorithm for 10,000 cycles, followed by conjugate gradient algorithm to relax the water molecules, counter ions and lipid tails while keeping the protein fixed with harmonic position restraints of 10 kcal.mol^-1^.Å^-2^ and lipid headgroups with 2.5 kcal.mol^-1^.Å^-2^. Next, the systems were then heated up from 10 to 310 K in the canonical ensemble (NVT) over 2 ns and subsequently maintained at 310 K for an additional 10 ns (divided into five 2 ns steps). During these equilibration steps, position restraints were sequentially reduced for both the protein (10 5 2.5 1 0.5 0.1 kcal.mol^-1^.Å^-2^) and the lipid headgroups (2.5 1 0.5 0.1 kcal.mol^-1^.Å^-2^).

Temperature control (310 K) was performed via Langevin dynamics ^90^, with a collision frequency of γ = 5.0 ps^-1^ during equilibration. Semi-isotropic position scaling with constant surface tension in the xy plane was applied to the protein-membrane systems. Pressure was controlled by coupling the system to a Berendsen barostat ^91^, at a reference pressure of 0.987 atm with a relaxation time of 1 ps. All equilibration runs have been carried out using an integration time step of 2 fs and a nonbonded cutoff of 10 Å. Hydrogen atoms were added assuming standard bond lengths and were constrained to their equilibrium position with the SHAKE algorithm. The final structures from the equilibration runs served as the starting point for the subsequent GaMD simulations.

#### WT and mutant domains

ColabFold PDB file was processed, solvated and neutralized as described above with a buffer of 25 Å. Minimization and equilibration protocol was updated as follows: during the second minimization step the prepped system was minimized using steepest descent algorithm for 10,000 cycles followed by conjugate gradient algorithm to relax the complete system. The system was then heated up from 10 to 310 K in the canonical ensemble (NVT) over 4 ns and subsequently maintained at 310 K for an additional 6 ns. After 10 ns of equilibration all restraints were released and ∼100 ns production runs were carried out in the isothermal-isobaric ensemble (NPT) for each system. The final structures from the production run served as the starting point for the subsequent GaMD simulations.

#### Gaussian accelerated MD (GaMD) simulations

System threshold energy was set to E = V_min_ + (V_max_ – V_min_)/k_0_ for all GaMD runs (iE = 2). Simulations were performed using a dual boost potential scheme, applying boosts to both the dihedral and total potential energy terms (igamd = 3). Each simulation proceeded through five sequential stages. First, a conventional MD preparatory phase of 0.4 ns was performed to equilibrate the system (ntcmdprep = 200,000). This was followed by conventional MD for 1.6 ns where maximum, minimum, average and standard deviation values of system potential (V_max_, V_min_, V_avg_ and *σ_v_*) were collected (ntcmd = 1,000,000). The system then entered GaMD preparatory equilibration phase of 0.4 ns where the boost potential was applied (ntebprep = 200,000). Next the GaMD equilibration was performed for 1.6 ns where potential statistics and boost parameters are updated every 0.1 ns (nteb = 1,000,000, ntave = 50,000). Finally, ∼500 ns or ∼1000 ns of GaMD simulations have been carried out in the NPT ensemble where the boost potential was applied and the boost parameters are fixed (nstlim = 251,000,000 or 502,000,000). The upper limit of the standard deviation of the first and second boost potential boost was set to 6 kcal/mol (sigma0D = sigma0P = 6.0). All GaMD simulations were conducted in triplicate using an integration time step of 2 fs. All GaMD simulations were performed using pmemd.cuda module of AMBER 20 software suite. A total of ∼25 µs of molecular simulations were carried out (∼0.7 µs of conventional MD and ∼24.5 µs of GaMD, see details in Table S3).

#### Energetic reweighting of GaMD simulations

Potential of mean force (PMF) profiles were calculated by reweighting the simulations using a 10^th^ order Maclaurin series expansion, as implemented in PyReweighting-2D.py ^47^. PMF calculations were performed per replica per system at 310 K, using root-mean-square deviation (RMSD, RC1) and radius of gyration (RoG, RC2) as reaction coordinates, computed using CPPTRAJ (check Structural analysis section for additional details). RC1/RC2 bins present across all replicas were then retained, and the corresponding PMF values were averaged and plotted for each system.

#### Structural Analysis

CPPTRAJ v6.18.1 was used to process and analyze the raw GaMD trajectories ^96^.

*Root-mean-square deviation (RMSD)* of Cα atoms was calculated using the final frame of the equilibration/production run as the reference structure, according to 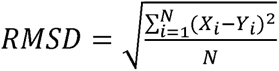, where N is the number of atoms, X_i_ is the coordinate vector of target atom i and Y_i_ is the coordinate vector for reference atom i.

*Root-mean-square fluctuations (RMSF)* of Cα atoms was calculated following removal of rotational and translational motions by aligning each frame to the initial frame of the production run, according to 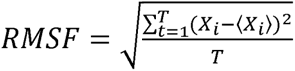, where X is the position of atom i at frame t, X is the average position of atom i and T is the total number of frames.

*Solvent accessible Surface Area (SASA)* was calculated for WT ANKLE2 (residues 1-235) and R236X using the LCPO algorithm ^97^.

*TM Angular kink* was defined as the angle formed between three points along the TM domain (2-4 Cα, 13-15 Cα, 32-34 Cα). *End-to-end distance* was defined as the distance between two points within the TM domain (1-2 Cα, 33-34 Cα).

*Salt bridge analysis* (LEM: R99-E103, CD: R215-E220, Unc1: E493-R536 and E533-R536) was performed by computing the distance between the oxygens atoms of negatively charged animo acid and nitrogen atoms of positively charged amino acid, the shortest distance between the N-O atoms was plotted to evaluate the salt bridge using a distance cutoff < 4 Å.

*Radius of Gyration (RoG)* of Cα atoms was calculated according to 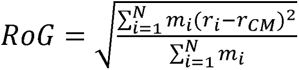, where N is the number of atoms, m_i_ is the mass of atom i, r_i_ is the position of atom i and r_CM_ is the center of mass.

Secondary structure propensities were assigned using the *DSSP* algorithm of Kabsch and Sander ^98^, which assigns secondary structure types for residues based on backbone amide and carbonyl atom positions.

*Interhelical distance* (WT CD, G201W and V229G) was calculated as the distance between the center of mass of α1 (residues 222-231) and α2 (residues 243-250) Cα atoms. Structural metrics were computed across all replicas and subsequently plotted using matplotlib and/or seaborn using a custom Python (v3.11.5) script.

#### Binary intramolecular GaMD contact analysis

Raw GaMD trajectories were processed and analyzed using MDAnalysis v2.9.0 ^99^. Each GaMD replica trajectory was first subsampled by extracting every 50^th^ frame. Intramolecular distances were then calculated for all atom pairs belonging to unique (non-identical) residues. Contact was defined as any atom pair with a distance of < 4 Å. To prevent overweighting the larger side chains, multiple qualifying contact pairs between the same two residues were collapsed into a single contact (e.g. if 5 atom pairs between residue i and residue j fell within 4 Å, this was counted as one contact). This presence/absence contact measure is termed as a “binary contact”. Raw binary contacts were reweighted using 10^th^ order Maclaurin series.

Reweighted binary contacts were calculated across all replicas for both WT and mutant systems, the mutant frequencies were subtracted from the WT frequencies (Δf = WT - Mutant) to obtain reweighted contact frequency difference. Residue-residue pairs with |Δf| > 0.34 were plotted as a waterfall plot using a custom Python (v3.11.5) script to identify reproducible changes in reweighted contact frequency between WT and mutant.

#### GaMD hydrogen bond analysis

Analysis was performed using MDAnalysis v2.9.0, with each GaMD replica trajectory subsampled by extracting every 50^th^ frame, consistent with the binary contact analysis above. Hydrogen bonds were identified across the protein backbone using a donor-acceptor distance cutoff of ≤ 3 Å and a donor-hydrogen-acceptor cutoff of ≥ 150°. Raw hydrogen bond frequencies were reweighted using 10^th^ order Maclaurin series.

Reweighted hydrogen bond frequencies were calculated across all replicas for both WT and mutant systems, and bonds exhibiting a > 10% reweighted frequency change relative to WT were plotted as a bar plot using a custom Python (v3.11.5) script.

#### Principal component analysis (PCA)

PCA was performed to capture the essential motions of the simulated systems. Covariance matrix of the protein backbone (N, Cα, C, O) atoms is calculated and diagonalized to obtain eigenvectors to describe the system motions. Each eigenvector (Principal Component (PC)) is sorted according to their eigenvalues, the first Principal Component (PC1) corresponds to the direction that maximizes variance of the system. GaMD trajectories in this work are projected into the collective coordinate space defined by the first two eigenvectors/PCs (PC1 and PC2), enabling the comparison between different ANKLE2 systems. To understand the differences between WT and mutant, backbone atoms of each simulated system was superposed and aligned to the same reference structure, allowing the projection onto the same collective coordinate space. PCA was performed using GROMACS 4.5.5 suite ^100^ (using g_covar and g_anaeig program).

#### Dynamic cross-correlation analysis (DCC_ij_)

DCC_ij_ analysis was performed to calculate Pearson’s correlations coefficients between the fluctuations of Cα atoms relative to their average positions ^42^. Dynamic cross-correlation coefficients (DCC_ij_) were calculated across three independent GaMD replicas, according to 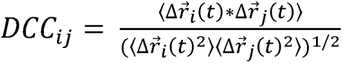, where Δr_i_ and Δr_j_ are the fluctuation vectors of atoms i and j respectively. The angle brackets denote the time average over the sampled simulation period. DCC_ij_ values range from −1 to 1, with positive values indicating correlated motion between atoms i and j, while negative values describing anticorrelated motion.

To enable comparison of inter-domain motion between WT ANKLE2 and R236X simulations, cross-domain DCC_ij_ values were block-averaged over six structural domains of ANKLE2 (TM: 1-34, LEM: 65-117, CD: 197-252, ANK: 299-472, Unc1: 473-620, Unc2: 824-876). This block-averaging analysis was performed over the course of the GaMD simulation and across all replicas.

### ANKLE2 domain recombinant protein production and purification

Plasmids were expressed in *E. coli* BL21* cells and grown in Terrific Broth at 37C and 225 rpm. When cultures reached OD_600_ between 2-3, expression was induced with 1 mM Isopropyl β-D-1-thiogalactopyranoside (IPTG, GoldBio) and performed overnight at 16°C. Cells were pelleted by centrifugation and resuspended in wash buffer (10% glycerol, 25 mM Tris, 30 mM imidazole, 250 mM NaCl, with a pH 8 of for WT CD or pH 6.8 for WT LEM) supplemented with lysozyme, DNAseI, 1 mM phenylmethylsulfonyl fluoride (PMSF) and a protease inhibitor cocktail (Thermo Scientific). Cells were lysed by sonication (Branson Analog Sonifier 250) for 3 x 2 min pulses, 40% Duty Cycle with a wide probe, incorporating a 2 min rest interval between each sonication. Lysates were clarified by ultracentrifugation at 35,000 *g* for 1 hour in Type 45 TI fixed-angle rotor (Beckman Coulter). The supernatants were passed onto a gravity column loaded with equilibrated Ni-NTA Agarose resin (Qiagen). The column was washed with the wash buffer and bound proteins were eluted with a gradient of 30 mM to 400 mM imidazole. Protein-containing fractions were pooled and the SUMO tag was removed by the addition of yeast His6-Ulp1 SUMO protease (purified in-house) during overnight dialysis at 4°C into wash buffer supplemented with 1 mM DTT. Samples were passed onto a fresh gravity column loaded with equilibrated Ni-NTA Agarose resin (Qiagen) and flowthrough was collected to isolate the untagged peptide. A second dialysis was performed overnight at 4°C into storage buffer (5% glycerol, 25 mM Tris, 100 mM NaCl, 0.5 mM DTT, with a pH 8 of for WT CD or pH 6.8 for WT LEM). Peptides were filtered and then concentrated using 2 kDa MWCO centrifugal filters (Cobetter).

### Circular dichroism spectroscopy

Samples were prepared by diluting concentrated protein to 30 μM in circular dichroism buffer (10 mM NaF, 25 mM phosphate, pH 7). Circular dichroism spectroscopy was performed at room temperature in a Jasco J-715 spectrometer with a scanning speed of 20 nm/min using a 1 mm pathlength quartz cuvette. Circular dichroism buffer was used as a blank, and the buffer spectrum was subtracted before analysis. Secondary structure was analyzed using BeStSel and the mean percent secondary structure across two independent measurements was plotted using GraphPad Prism 11 ^101^.

### Data visualization

Figures were assembled using Affinity Designer (v2.6.5). Some graphics were made using BioRender. Microscopy images, blots, and gels were prepared using Fiji/ImageJ (v1.54j) ^85^. Graphs were generated using either GraphPad Prism 11, RStudio (v2026.01.2), or Python (v3.11.5). Protein structures were evaluated using UCSF ChimeraX (v1.6.1)^94^. Phosphoproteomic kinase-substrate networks were generated using CytoScape (v3.10.2) ^84^.

