## Supplementary Information for "Integrative proteomic analysis and molecular dynamics simulations of ANKLE2 reveal mechanisms of microcephaly"

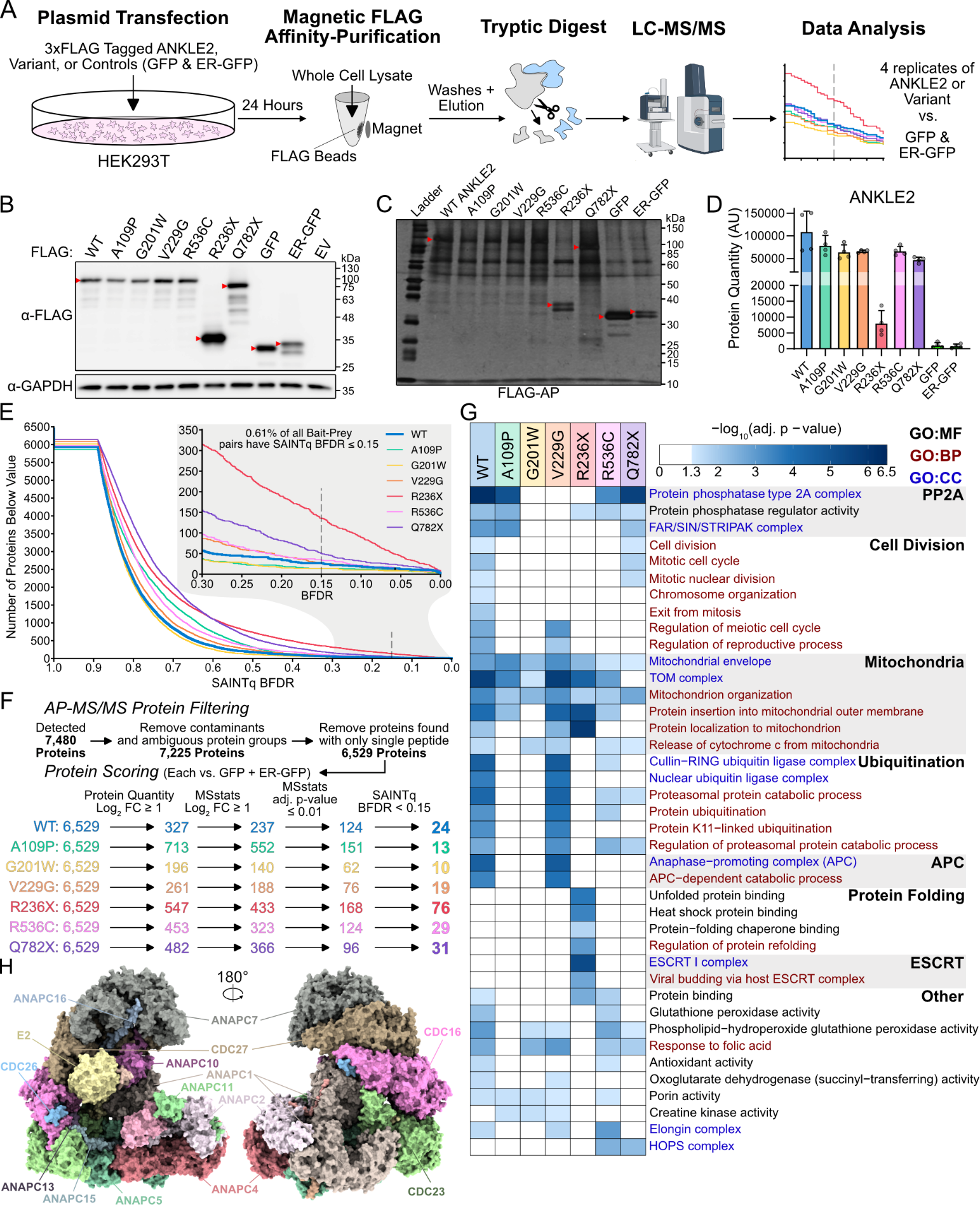
**Supplementary Information**

**Figure S1: Proteomic analysis of ANKLE2 and pathogenic variants. Related to Figures 1 and 2. A)** Schematic of proteomics pipeline starting with expression of ANKLE2 variants in HEK293T cells, followed by FLAG affinity-purification, tryptic digest, and tandem mass spectrometry. **B)** Representative western blot of ANKLE2 variant and control expression in HEK293T cells after transfection. **C)** Silver stain of affinity-purified samples show relative abundance of co-purified proteins. Red arrows indicate expected ANKLE2 (or variant) band. **D)** Raw protein quantities of ANKLE2 detected from proteomics. Gray circles indicate biological replicates. **E)** Waterfall plot of the number of proteins below SAINTq BFDR values. **F)** Flow chart for number of proteins that fulfill each criterion with final number of high-confidence interactions bolded. **G)** Functional enrichment of high-confidence interactors using g:Profiler ^28^ shows profound loss of protein-interactions involved in cell division for pathogenic variants. **H)** Labeled structure of anaphase-promoting complex (pdb: 5A31) ^36^.

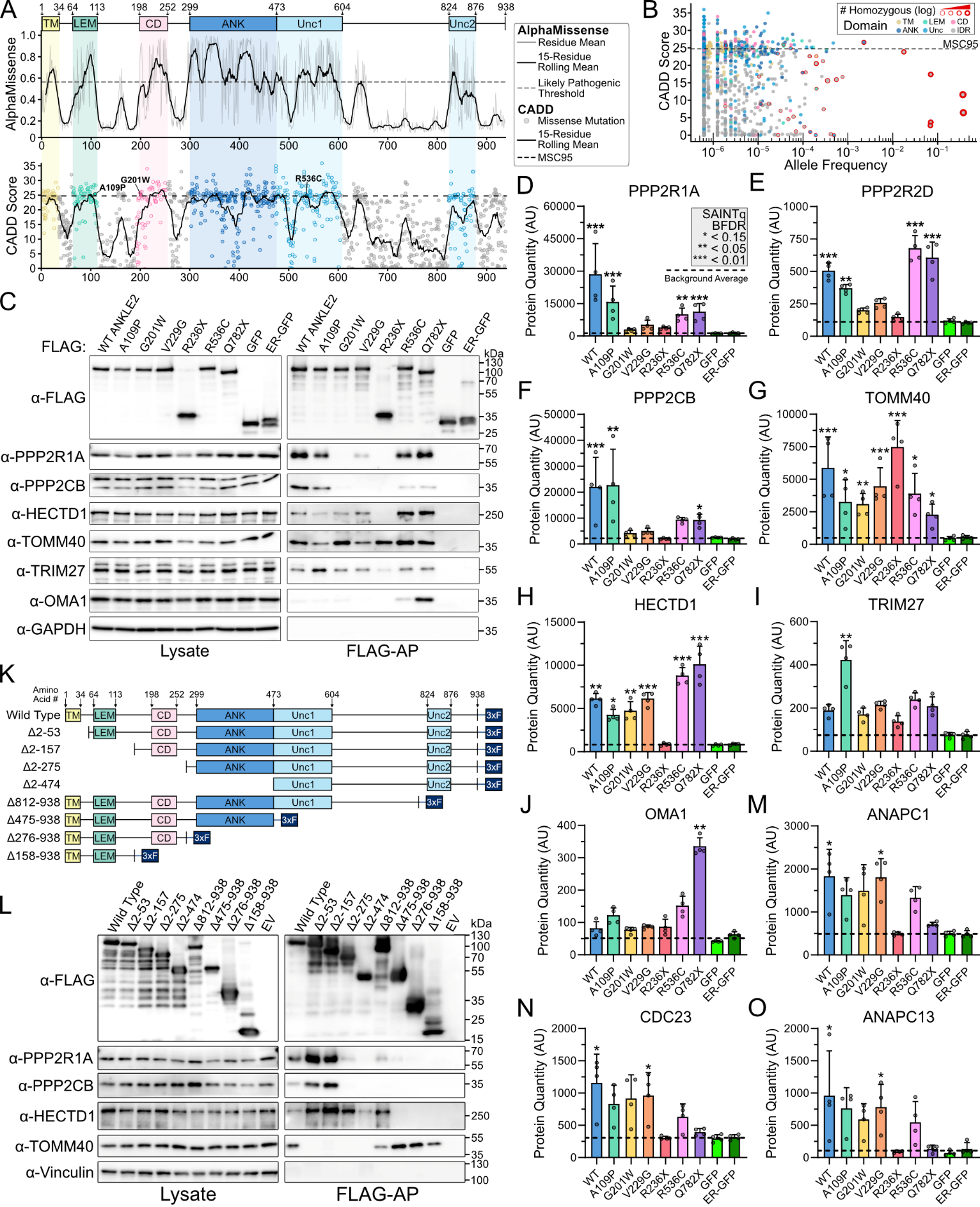

**Figure S2: Functional validation of ANKLE2 interactions. Related to Figure 2. A)** Alpha Missense ^103^ and CADD scores ^22^ plotted against ANKLE2 residue. **B)** CADD scores of missense mutations plotted against allele frequency from the gnomAD database ^22^. Dot color indicates domain and red border thickness indicates number of homozygous occurrences. **C)** Western blot validation of ANKLE2 interactions from proteomics screen. **D-J)** Raw protein quantity measurements from proteomics match trends in western blot validation. Dashed line indicates average of all eight control measurements (GFP and ER-GFP). Stars indicate SAINTq BFDR value. **K)** Schematic of ANKLE2 truncations used to identify regions mediating ANKLE2 interactions. **L)** Western blot of FLAG co-immunoprecipitation assay using ANKLE2 truncation interactions. **M-O)** Raw protein quantity measurements from proteomics of high-confidence APC members.

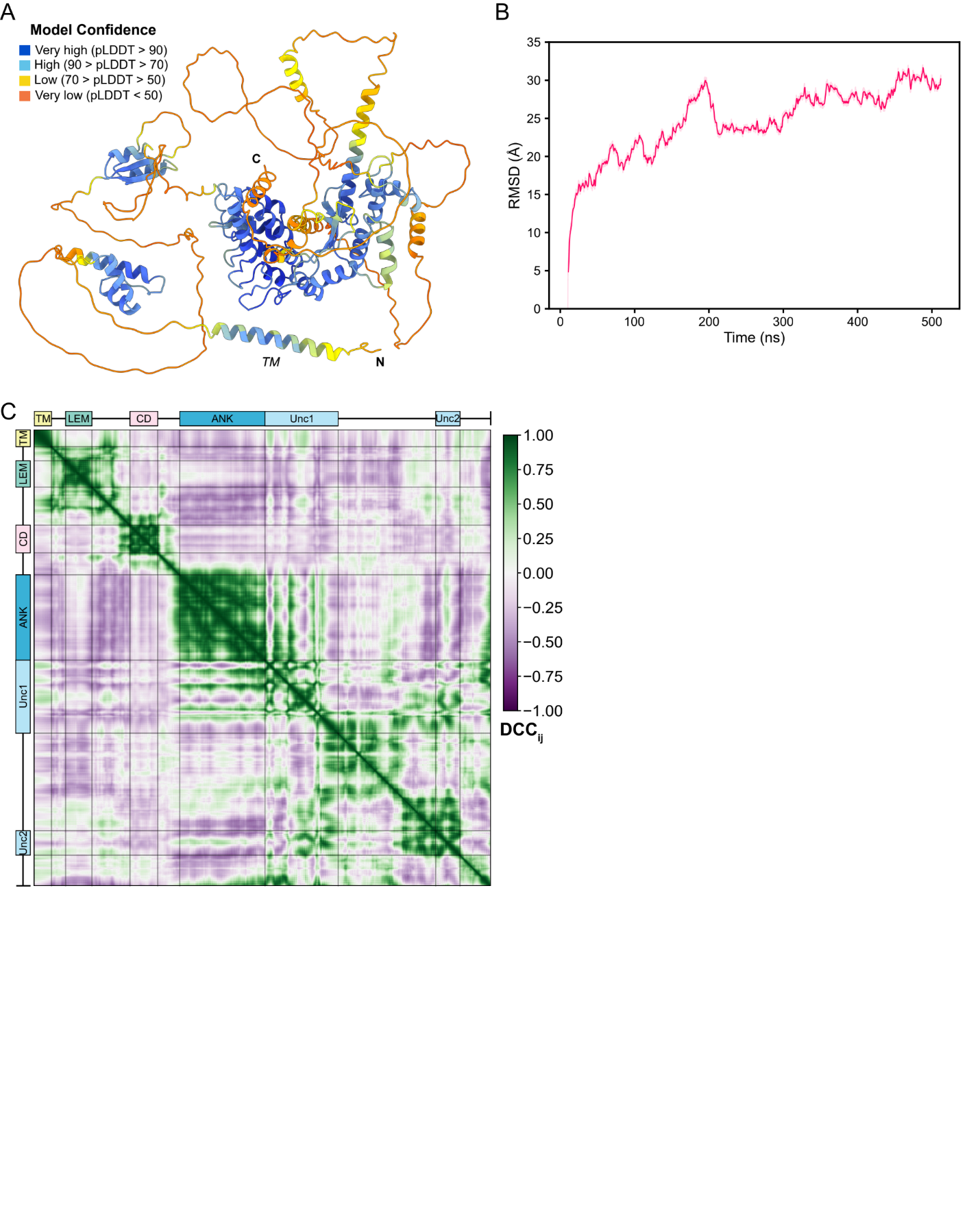

**Figure S3: System preparation and validation of WT ANKLE2 GaMD simulations. Related to Figure 3. A)** Ribbon representation of the ColabFold-predicted structure of ANKLE2 with TM domain, N- and C-termini annotated. Per-residue pLDDT scores are overlaid on the structure. **B)** root-mean-square deviation (RMSD) of ANKLE2-WT Δ1-34 Cα atoms relative to the equilibrated structure. Muted lines show raw RMSD values at each frame, solid lines show the rolling average (window = 1 ns). **C)** Dynamic cross-correlation (DCC_ij_) for Cα atoms of WT ANKLE2, calculated across all replicas. The replica-averaged DCC_ij_ is shown. Residues corresponding to structured domains are annotated.

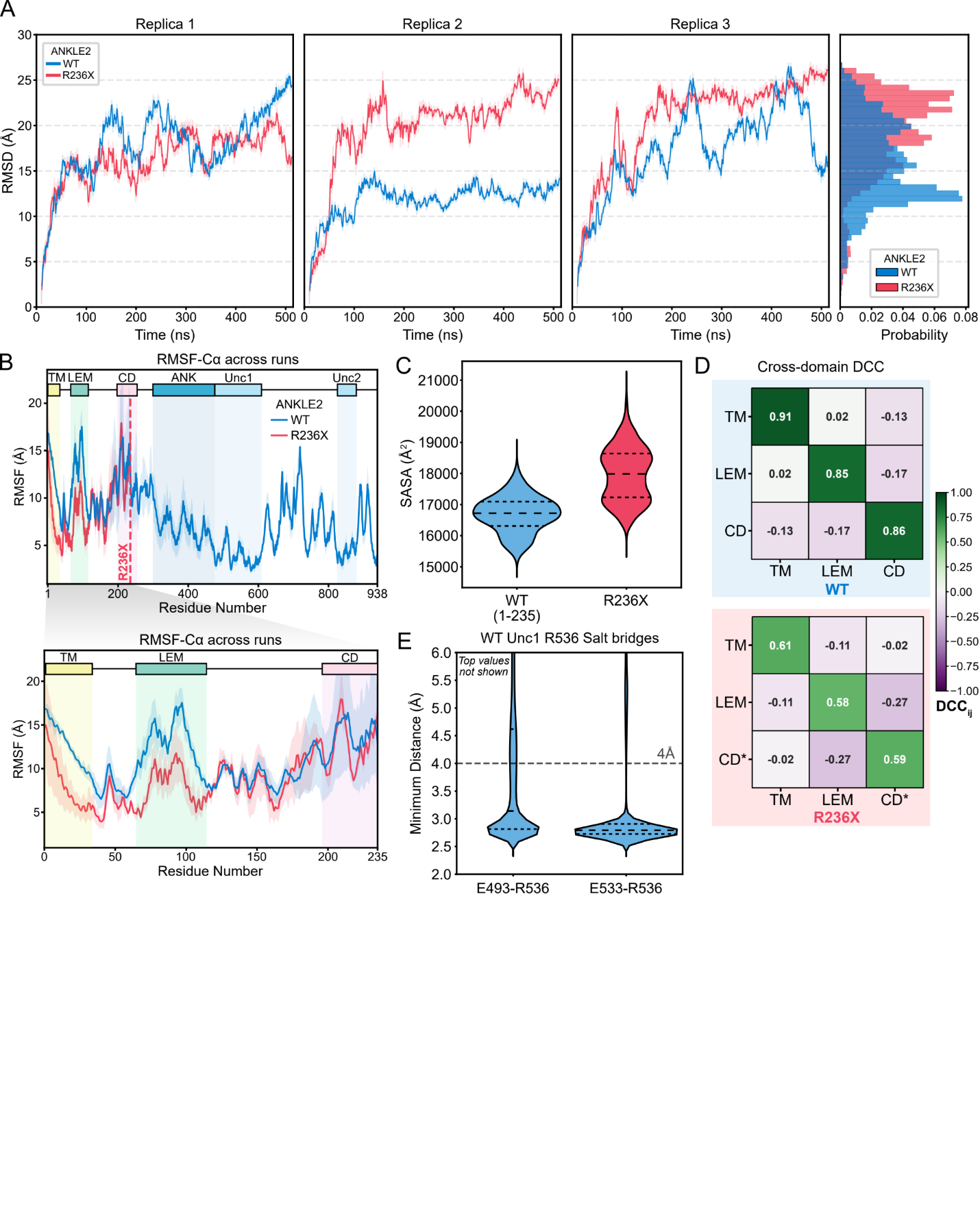

**Figure S4: GaMD simulations of nonsense mutation R236X. Related to Figure 4. A)** Per-replica Cα root-mean-square deviation (RMSD) for WT ANKLE2 (blue) and R236X (red), calculated relative to the structure at the end of the equilibration run. Muted lines show raw RMSD values at each frame, solid lines show the rolling average (window = 1 ns). (Right) Probability density of RMSD values pooled across replicas for each mutant. **B)** Root-mean-square fluctuation (RMSF) of Cα atoms for WT ANKLE2 and R236X across three independent replicas (Top). RMSF of Cα atoms for WT ANKLE2 (residues 1-235) and R236X across three independent replicas (Bottom). Solid lines indicate the mean RMSF, shaded regions indicate the standard deviation and residues corresponding to structured domains are annotated. **C)** Solvent accessible surface area (SASA) (Å^2^) of WT ANKLE2 (residues 1-235) and R236X, pooled across all replicas per variant and shown as a violin plot (dashed lines represent quartiles). **D)** Domain-averaged dynamic cross-correlation (DCC_ij_) among WT ANKLE2 and R236X structured domains (TM, LEM, CD), calculated for Cα atoms and block averaged to yield domain-specific correlation values. CD* represents truncated CD within R236X. **E)** Minimum distance between the nitrogen atoms of R536 and oxygen atoms of E493/E533, pooled across all replicas per variant and shown as a violin plot (dashed lines represent quartiles). The salt bridge distance cutoff (4 Å) is indicated.

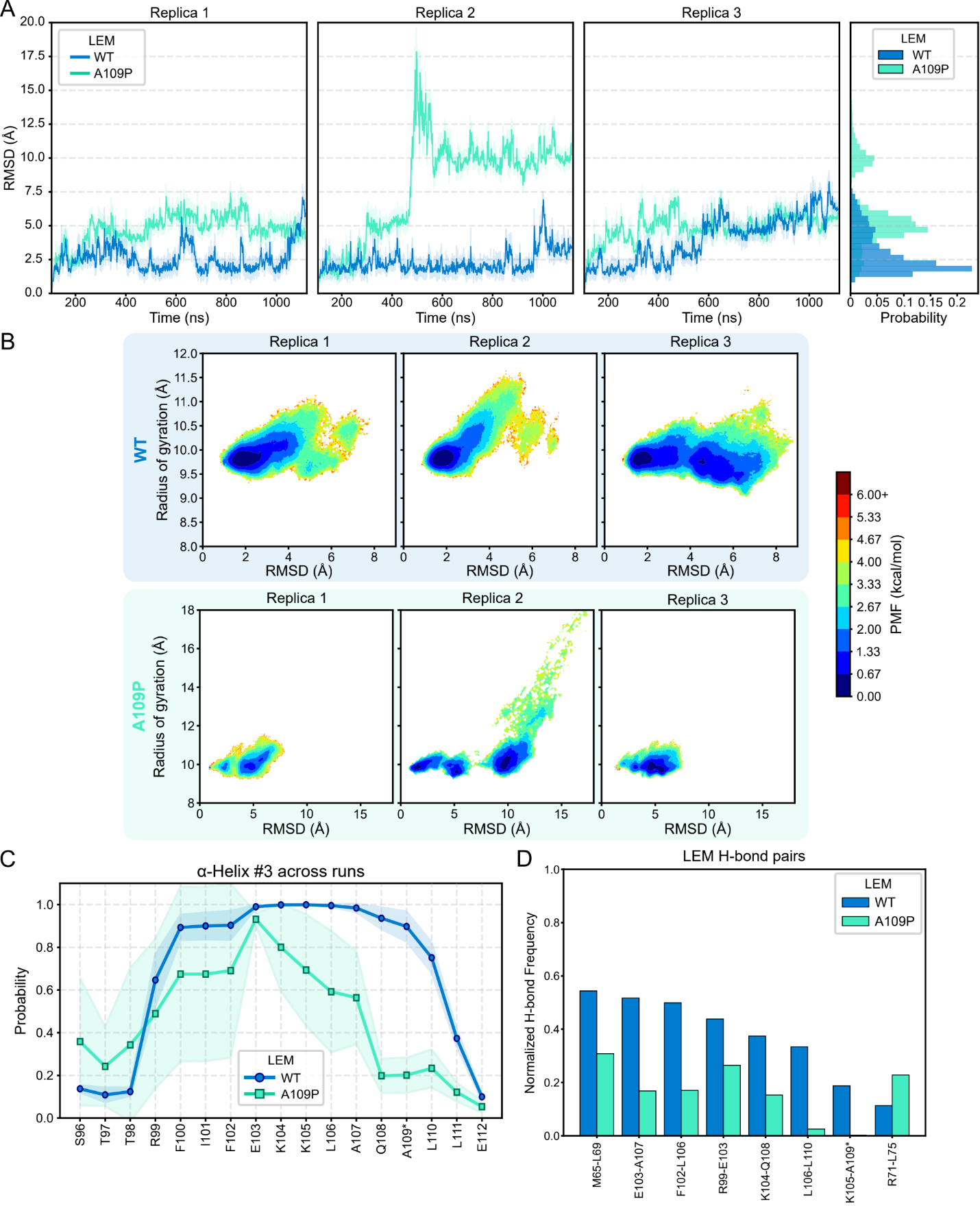

**Figure S5: GaMD simulations reveal A109P-induced destabilization of the LEM domain. Related to Figure 5. A)** Per-replica Cα root-mean-square deviation (RMSD) for WT LEM (blue) and A109P (green), calculated relative to the structure at the end of the production run. Muted lines show raw RMSD values at each frame, solid lines show the rolling average (window = 1 ns). (Right) Probability density of RMSD values pooled across replicas for each mutant. **B)** 2D potential mean force (PMF) per replica landscape for WT LEM (Top) and A109P (Bottom) with RMSD and radius of gyration (RoG) as the reaction coordinates. **C)** DSSP-assigned secondary structure probability for α3, pooled across replicas for each mutant. **D)** Reweighted Hydrogen bond (H-bond) frequency of WT LEM and A109P, pooled across replicas. Residue pairs exhibiting a >10% change in reweighted frequency relative to WT are shown.

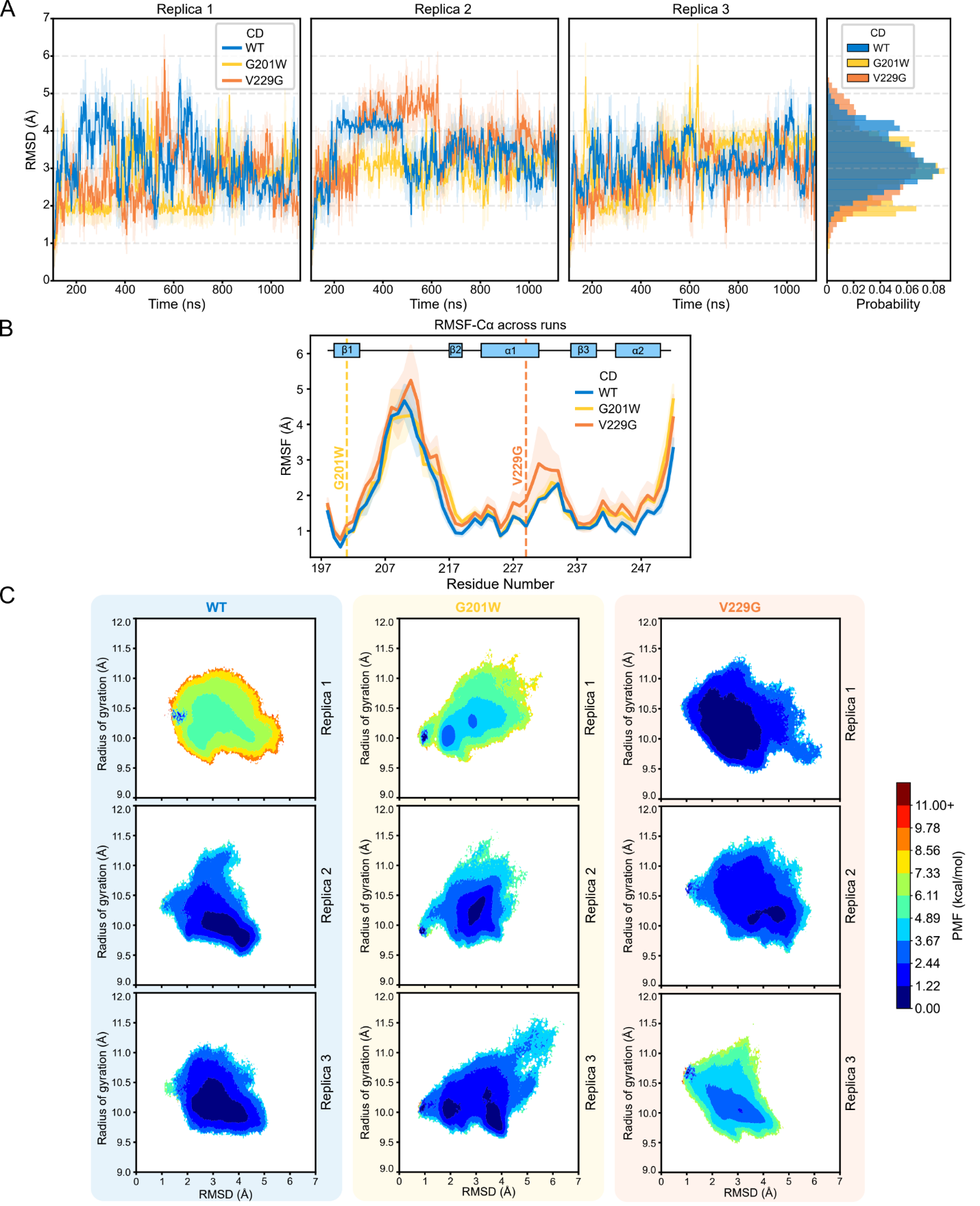

**Figure S6: Low-energy conformation states of CD mutants, G201W and V229G. Related to Figure 6. A)** Per-replica Cα root-mean-square deviation (RMSD) for WT CD (blue), G201W (yellow), and V229G (orange), calculated relative to the structure at the end of the production run. Muted lines show raw RMSD values at each frame, solid lines show the rolling average (window = 1 ns). (Right) Probability density of RMSD values pooled across replicas for each mutant. **B)** Root-mean-square fluctuation (RMSF) of Cα atoms for WT CD, G201W, and V229G across three independent replicas. Solid lines indicate the mean RMSF, shaded regions indicate the standard deviation. Residues corresponding to secondary structural elements and mutation site are annotated. **C)** 2D potential mean force (PMF) per replica landscape for WT CD (Top), G201W (Middle), and V229G (Bottom) with RMSD and radius of gyration (RoG) as the reaction coordinates.

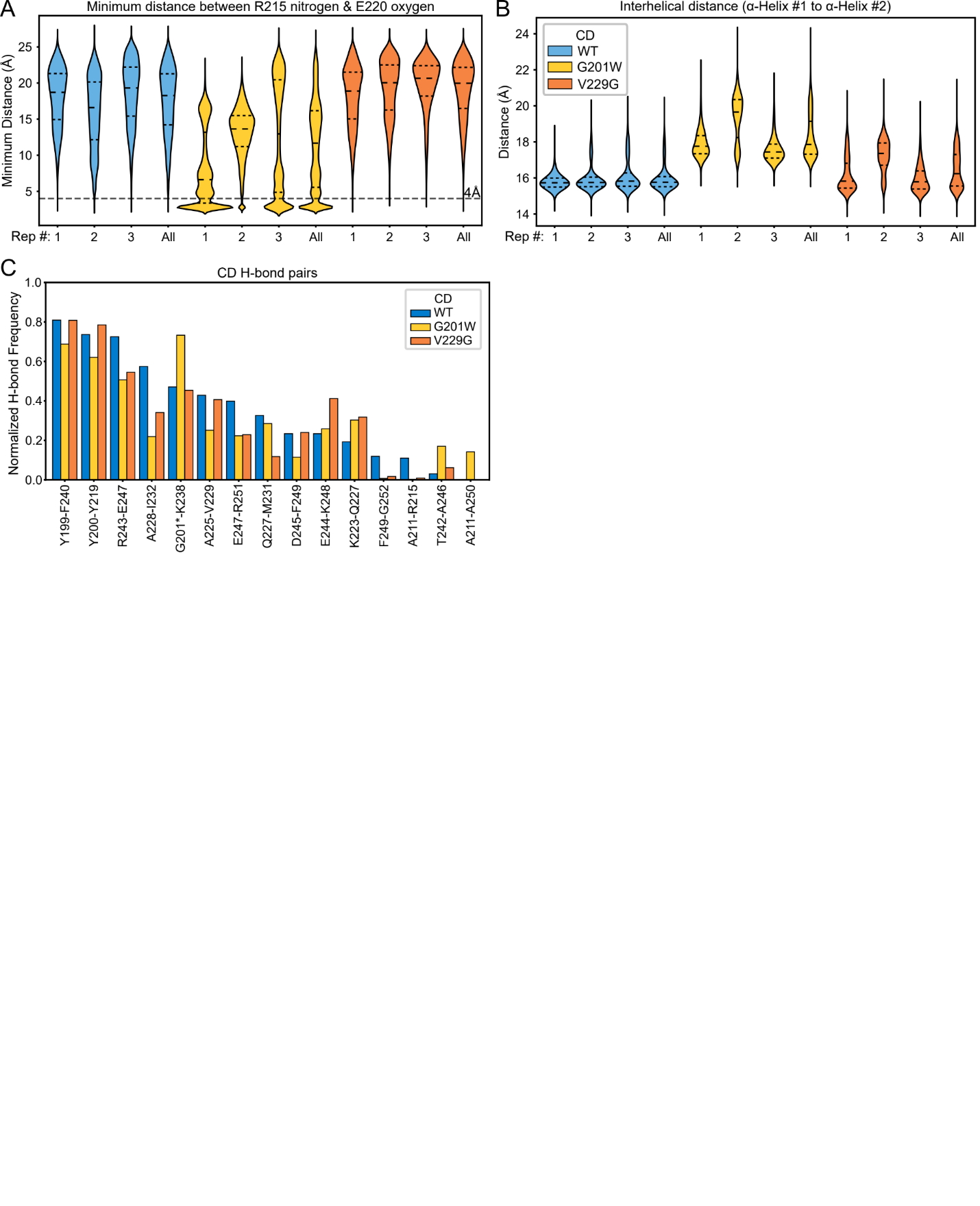

**Figure S7: GaMD simulations reveal structural changes in mutants with impaired PP2A binding. Related to Figure 6. A)** Minimum distance between the nitrogen atoms of R215 and oxygen atoms of E220, per replica and pooled across all replicas per mutant and shown as a violin plot. The salt bridge distance cutoff (4 Å) is indicated. **B)** Distance between the center of mass of the Cα atoms (representing the geometric center) of α1 and α2 within the CD domain, per replica and pooled across all replicas per mutant. **C)** Reweighted Hydrogen bond (H-bond) frequency of WT CD, G201W, and V229G, pooled across replicas. Residue pairs exhibiting a >10% change in reweighted frequency relative to WT are shown.

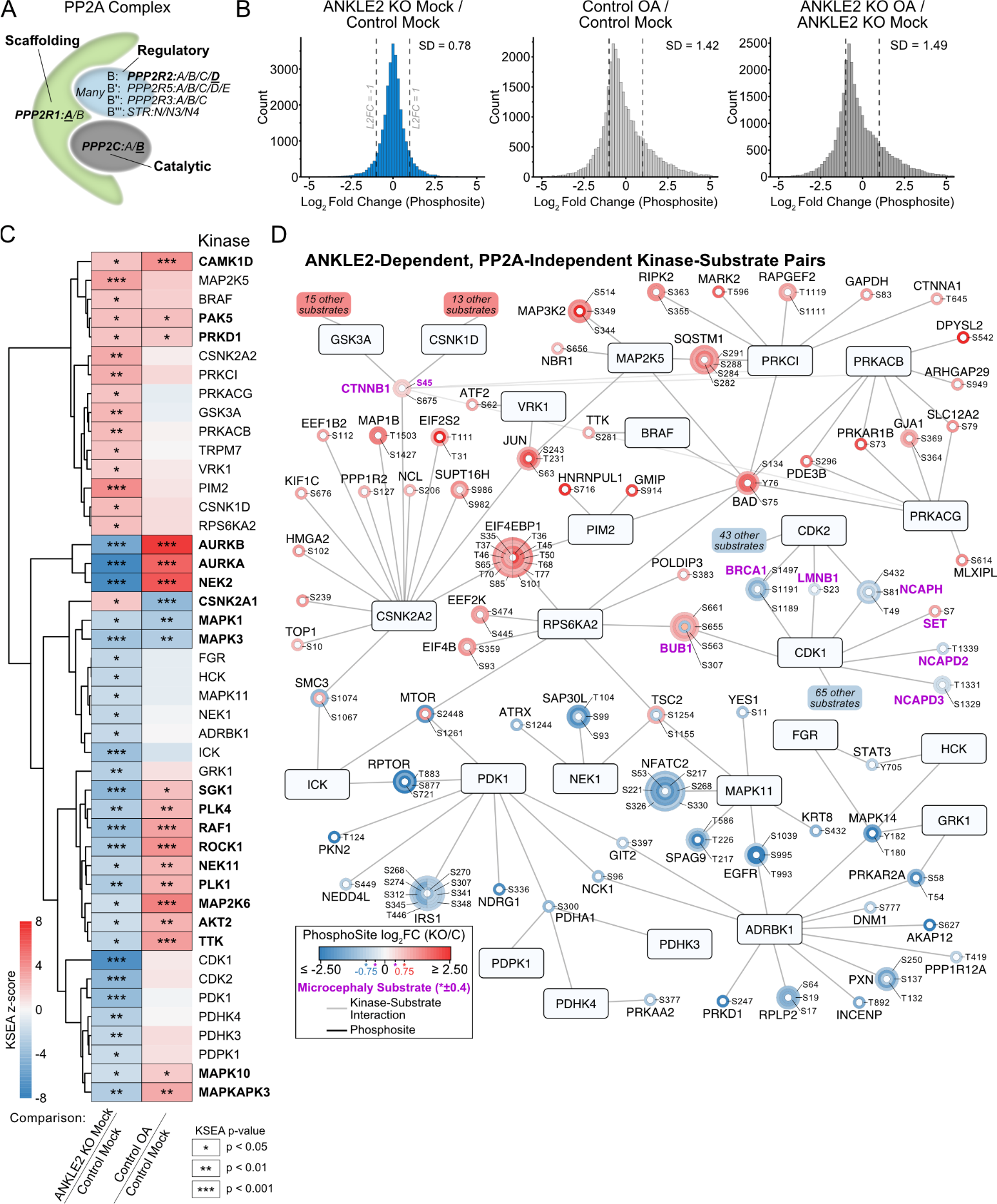

**Figure S8: PP2A-dependent versus -independent regulation of the phosphoproteome by ANKLE2. Related to Figure 7. A)** Schematic of the PP2A complex subunits. Subunits that interact with ANKLE2 (PPP2R1A, PPP2CB, and PPP2R2D) are bolded. **B)** Histograms of all phosphosite fold changes for mock treated ANKLE2 KO cells or OA treated controls cells versus mock control. **C)** Full list of significant kinases from KSEA of ANKLE2 KO-Mock vs. Control-Mock which indicate ANKLE2-dependent changes. Significance values for the same kinases in Control-OA vs. Control-Mock were used to differentiate PP2A-dependent from PP2A-independent. **D)** Kinase-substrate network of ANKLE2-dependent and PP2A-independent kinases with matching directional substrate |log_2_ fold changes| ≥ 0.75. Microcephaly substrates (pink) with |log_2_ fold changes| ≥ 0.40 are also shown. Kinases associated with more than 200 identified phosphosites (m ≥ 150: CDK1, CDK2, GSK3A, CSNK1D) were limited to only microcephaly substrates. The number of these hidden substrate proteins, irrespective of number of phosphosites or other kinase interactions, is indicated in colored boxes that correspond to kinase z-score.

**Supplementary Data 1: ANKLE2 AP-MS proteomic data of ANKLE2 and g:Profiler enrichment analysis.**

**Supplementary Data 2: ANKLE2 domain circular dichroism.**

**Supplementary Data 3: Phosphoproteomic data of Huh7 ANKLE2 KO cells.**

**Supplementary Video 1: Molecular dynamics simulation of WT and A109P LEM domain backbone H-bond changes**

**Supplementary Video 2: Molecular dynamics simulation of WT and G201W CD domain salt bridge formation**

**Table S1: Curated list of microcephaly genes. Related to Figure 6.**

| **Gene name** | **UniProt ID** | **Synonyms** | **Reference PMID(s)** (First 3) |
| --- | --- | --- | --- |
| AGMO | Q6ZNB7 | TMEM195 | 27000257 |
| AKT3 | Q9Y243 | PKBG | 31929334, 32827175, 40032969 |
| ANKLE2 | Q86XL3 | LEM4, MCPH16 | 25259927, 30214071, 31735666 |
| AP4E1 | Q9UPM8 |  | 20972249, 36226339 |
| APOE | P02649 |  | 17272620, 20002130 |
| ARFGEF2 | Q9Y6D5 | BIG2 | 12682315, 14647276, 23812912 |
| ASPM | Q8IZT6 | MCPH5 | 12355089, 14574646, 16141009 |
| ATR | Q13535 | FRP1 | 12640452, 17564965 |
| BRCA1 | P38398 | RNF53 | 32843487, 38146508, 40519302 |
| BUB1 | O43683 | BUB1L, MCPH30 | 35044816 |
| CASK | O14936 | LIN2 | 19165920, 21954287, 25886057 |
| CDK4 | P11802 | MCPH31 | 40210435 |
| CDK5RAP2 | Q96SN8 | CEP215, MCPH3 | 10677332, 15793586, 23995685 |
| CDK6 | Q00534 | CDKN6, MCPH12 | 23918663 |
| CENPE | Q02224 | MCPH13 | 24748105 |
| CEP135 | Q66GS9 | CEP4, MCPH8 | 22521416, 26657937 |
| CEP152 | O94986 | MCPH9 | 15806441, 20598275, 22775483 |
| CEP295 | Q9C0D2 | KIAA1731 | 38154379 |
| CEP63 | Q96MT8 |  | 21983783 |
| CHEK1 | O14757 | CHK1 | 16217032, 19546241, 36549593 |
| CIT | O14578 | CRIK, MCPH17, STK21 | 27453578, 27453579, 27503289 |
| COPB2 | P35606 | MCPH19 | 29036432, 34450031 |
| COX20 | Q5RI15 | FAM36A | 30656193, 32606554 |
| CPAP | Q9HC77 | CENPJ, LAP, LIP1, MCPH6 | 12843329, 16900296, 20978018 |
| CTNNB1 | P35222 | CTNNB, NEDSDV | 25326669, 27915094, 23033978 |
| CTU2 | Q2VPK5 | MFRG, NCS2 | 27480277, 31301155 |
| DEAF1 | O75398 | SPN, ZMYND5 | 24668509, 26834045 |
| DYRK1A | Q13627 | DYRK, MNB, MNBH | 18405873, 21294719, 23160955 |
| EFTUD2 | Q15029 | SNRP116 | 22305528, 22541558, 23188108 |
| EIF2S3 | P41091 | EIF2G | 27333055 |
| EOMES | O95936 | TBR2 | 17353897, 18794345 |
| FEZF2 | Q8TBJ5 | FEZL, ZNF312 | 21471212, 34562292 |
| FOXG1 | P55316 | FKH2, FKHL1, FKHL2, FKHL3, FKHL4, FOXG1A, FOXG1B, FOXG1C | 16133170, 18571142, 21441262 |
| HMGB3 | O15347 | HMG2A, HMG4 | 4998085, 24993872 |
| HNRNPU | Q00839 | HNRPU, SAFA, U21.1 | 25356899 |
| IGF1 | P05019 | IBP1, IGF-1 | 21396584, 38838658, 38952118 |
| IGF1R | P08069 | IGF1RES | 22130793, 23045302, 26252249 |
| IGFALS | P35858 | ALS | 38838658 |
| KIF11 | P52732 | EG5, KNSL1, TRIP5 | 22284827, 26472404, 27212378 |
| KIF14 | Q15058 | MCPH20 | 28892560, 23308235, 28892560 |
| KIF20B | Q96Q89 | KRMP1, MPHOSPH1 |  |
| KNL1 | Q8NG31 | CASC5, MCPH4 | 10521316, 26626498, 27149178 |
| LAGE3 | Q14657 | DXS9879E, ESO3, ITBA2 | 28805828 |
| LMNB1 | P20700 | LMN2, LMNB, MCPH26 | 32910914, 33033404 |
| LMNB2 | Q03252 | LMN2, MCPH27 | 33033404, 40011009 |
| MBD5 | Q9P267 | KIAA1461 | 19904302, 21271666, 33912662 |
| MCPH1 | Q8NEM0 |  | 12046007, 11857108, 15199523 |
| MECP2 | P51608 |  | 15557528, 12481990 |
| MFSD2A | Q8NA29 | MFSD2, NLS1 | 24828044, 26005868,26005865 |
| MRE11A | P49959 | HNGS1, MRE11 | 15574463, 21227757 |
| MRTFB | Q9ULH7 | KIAA1243, MKL2 | 23692340, 25755095 |
| MSI1 | O22467 |  | 28572454 |
| MSMO1 | Q15800 | DESP4, ERG25, SC4MOL | 21285510, 24144731 |
| NCAPD2 | Q15021 | CAPD2, CNAP1, KIAA0159, MCPH21 | 27737959, 28097321, 31056748 |
| NCAPD3 | P42695 | CAPD3, KIAA0056, MCPH22 | 27737959 |
| NCAPH | Q15003 | BRRN, BRRN1, CAPH, KIAA0074, MCPH23 | 27737959 |
| NDE1 | Q9NXR1 | NUDE | 21529752, 22526350, 30637988 |
| NSD1 | Q96L73 | ARA267, KMT3B | 16770806, 19844260, 21567906 |
| NUP37 | Q8NFH4 | MCPH24 | 30179222 |
| OSGEP | Q9NPF4 | GCPL1 | 28805828 |
| PAX6 | P26367 | AN2 | 7951315, 17406642, 26130484 |
| PDCD6IP | Q8WUM4 | AIP1, ALIX, KIAA1375, MCPH29 | 32286682 |
| PHC1 | P78364 | EDR1, PH1, MCPH11 | 23418308 |
| PLK4 | O00444 | SAK, STK18, MCCRP2 | 25344692, 25320347 |
| PNKP | Q96T60 | MCSZ | 20118933, 23224214 |
| POMT1 | Q9Y6A1 |  | 19299310, 17878207 |
| POMT2 | Q9UKY4 |  | 19299310, 17878207,17634419 |
| PPIL1 | Q9Y3C6 | CYPL1 | 33220177 |
| PQBP1 | O60828 | NPW38 | 15024694, 6711604,14634649 |
| PRSS12 | P56730 | MRT1, NETR | 13636700, 13738004 |
| PRUNE1 | Q86TP1 | PRUNE, NMIHBA | 28334956, 26539891 |
| RRP7A | Q9Y3A4 | MCPH28 | 33199730 |
| RTTN | Q86VV8 | MSSP | 22939636, 26608784, 26940245 |
| SASS6 | Q6UVJ0 | SAS6, MCPH14 | 24951542, 30639237 |
| SEL1L | Q9UBV2 | TSA305 | 37943610, 37943617, 41862645 |
| SET | Q01105 |  | 21515671, 29688601 |
| SLC25A19 | Q9HC21 | DNC 1, MUP1, MCPHA | 12185364 |
| SLC2A1 | P11166 | GLUT1 | 20687207, 18606970 |
| SNX3 | O60493 |  | 12471201 |
| STIL | Q15468 | SIL, MCPH7 | 22989186, 19215732, 25218063 |
| SYVN1 | Q86TM6 | HRD1, KIAA1810 | 37943610 |
| TAF2 | Q6P1X5 | CIF150, TAF2B | 24084144, 22633631 |
| TBX2 | Q13207 |  | 20206336, 21271665, 22052739 |
| TBX4 | P57082 |  | 20206336, 21271665, 22052739 |
| TP53RK | Q96S44 | PRPK | 28805828 |
| TPRKB | Q9Y3C4 | CGI-121, My019 | 28805828 |
| TRA2B | P62995 | SFRS10 | 36549593 |
| TRAIP | Q9BWF2 | RNF206, TRIP SCKL9 | 26595769,11424145, 34235748 |
| TRAPPC14 | Q8WVR3 | C7orf43, MAP11, MCPH25 | 30715179 |
| TRAPPC9 | Q96Q05 | KIAA1882, NIBP | 20004763, 20004765, 21629298 |
| TRMT10A | Q8TBZ6 | RG9MTD2, MSSGM1 | 24204302, 25053765 |
| TSEN15 | Q8WW01 | PCH2F, SEN15 | 25558065, 27392077 |

**Table S2: Antibodies. Related to STAR Methods.**

| **Antibody** | **Host Species** | **Dilution Used** | **Supplier (Catalog #)** | **RRID** |
| --- | --- | --- | --- | --- |
| GAPDH | Mouse | 1:1000 | Fisher (MA-15738) | AB_2537652 |
| FLAG | Mouse | 1:1000 | MilliporeSigma (F1804) | AB_262044 |
| ANKLE2 | Rabbit | 1:1000 | Bethyl Labs (A302-966A-M) | AB_2780882 |
| PPP2R1A | Rabbit | 1:1000 | Abcam (ab168371) | AB_2892220 |
| PPP2CB | Rabbit | 1:1000 | Abcam (ab154551) | N/A |
| HECTD1 | Rabbit | 1:1000 | Proteintech (20605-1-AP) | AB_10732804 |
| TOMM40 | Mouse | 1:1000 | SantaCruzBio (sc-365467) | AB_10847086 |
| TRIM27 | Rabbit | 1:1000 | Proteintech (12205-1-AP) | AB_2256660 |
| OMA1 | Rabbit | 1:1000 | Proteintech (17116-1-AP) | AB_2299053 |
| Vinculin | Mouse | 1:1000 | Sigma (V9131) | AB_477629 |
| P-GEFH1 (Ser886) | Rabbit | 1:1000 | CST (14143) | AB_2798402 |
| BAF | Rabbit | 1:1000 | Gift from Dr. Matthew Wiebe | N/A |
| TOMM20 | Rabbit | 1:1000 | Proteintech (11802-1-AP) | AB_2207530 |
| TFAM | Rabbit | 1:5000 | Proteintech (22586-1-AP) | AB_11182588 |
| DRP1 | Rabbit | 1:2000 | Proteintech (12957-1-AP) | AB_2093525 |
| COX IV | Rabbit | 1:1000 | Abcam (ab202554) | AB_2861351 |
| Deadpan | Rat | 1:1000 | Abcam (ab195173) | AB_2687586 |
| pHH3 | Rabbit | 1:1000 | MilliporeSigma (06-570) | AB_310177 |
| Anti-Rat-Alexafluor-647 | Donkey | 1:500 | Jackson ImmunoResearch Laboratories (712-605-153) | AB_2340694 |
| Anti-Rabbit-Alexafluor-488 | Donkey | 1:500 | Jackson ImmunoResearch Laboratories (711-545-152) | AB_2313584 |
| Anti-Mouse-HRP | Rabbit | 1:5000 | SouthernBiotech (6170-05) | AB_2796243 |
| Anti-Rabbit-HRP | Goat | 1:5000 | SouthernBiotech (4030-05) | AB_2687483 |

**Table S3: MD and GaMD simulations. Related to STAR Methods.**

| **System** | **# of Atoms** | **# of MD Simulations** | **# of GaMD Simulations** |
| --- | --- | --- | --- |
| WT LEM | ~58k | 1x 0.1 µs | 3x ~1 µs |
| A109P | ~58k | 1x 0.1 µs | 3x ~1 µs |
| WT CD | ~67k | 1x 0.1 µs | 3x ~1 µs |
| G201W | ~73k | 1x 0.1 µs | 3x ~1 µs |
| V229G | ~65k | 1x 0.1 µs | 3x ~1 µs |
| WT Unc1 | ~131k | 1x 0.1 µs | 3x ~1 µs |
| R536C | ~135k | 1x 0.1 µs | 3x ~1 µs |
| ANKLE2 Δ1-34 | ~598k | - | 1x ~0.5 µs |
| WT ANKLE2 | ~815k | - | 3x ~0.5 µs |
| R236X | ~559k | - | 3x ~0.5 µs |
